# From Perception to Appraisal: Brain Responses to Natural and Built Features in Urban Environments

**DOI:** 10.1101/2025.10.31.685763

**Authors:** Carolina Zähme, Isabelle Sander, Aleksandrs Koselevs, Simone Kühn, Klaus Gramann

## Abstract

Exposure to urban rather than natural environments has been linked to elevated stress and diminished well-being, prompting interest in the underlying mechanisms and influences of specific urban elements. This study investigated how semantic and lower-level visual characteristics of urban scenes influence subjective experience and underlying neural processes. A total of 63 adults viewed street-level photographs depicting urban environments with varying proportions of vegetation, buildings, vehicles, and sky visibility while electroencephalography (EEG) was recorded. Subjective ratings indicated that urban scenes with more greenery compared to built elements were consistently perceived as more positive. In terms of lower-level image features, images with higher non-straight edge density also led to scenes being perceived as more positive. In contrast, a more visible sky area and a higher mean metric depth led to more negative scene appraisal. On a neural level, urban scenes with higher greenery/built ratios enhanced visual evoked P1 amplitudes, while higher straight-edge density led to more pronounced N1 amplitudes. Further analysis showed that later P3 and LPP components were associated with subjective ratings on higher-level evaluative but not basic affective dimensions. These results partially replicate previous studies, suggesting that urban greenery exerts an effect on subjective well-being and influences early visual processing. Moving forward, Neuro-Urbanism research should employ more diversified stimuli and leverage immersive or in situ paradigms to refine our understanding of the relationship between greenery in urban environments and well-being to inform evidence-based urban design strategies.

**Highlights:**

- Higher proportions of greenery in urban environments lead to more positive responses across a variety of subjective rating scales.
- More urban greenery elicited more pronounced P1 amplitudes, replicating previous findings.
- Earliest visual processing (P1) responds to natural elements of the environment, before processing of visual features characteristic of artificial elements (N1).
- Later ERP components (P3, LPP) are associated with subjective scene appraisal of higher level and aesthetic evaluation, pointing to a separation between perceptual and evaluative processes.

## 1. Introduction

### 1.1. Associations between mental health and the living environment

Urban environments are increasingly recognized as powerful determinants of mental health. Today, more than half of the global population resides in cities, and this proportion is projected to rise substantially in the coming decades (United Nations, 2025) as cities offer advantages like dense infrastructure, education, and cultural life (Cohen, 2006). Yet, these benefits come with a high concentration of stressors such as traffic noise, air pollution, and heat-island effects (Feng et al., 2023; Moore et al., 2003). These ambient stressors converge with psychosocial pressures, including anonymity, weakened social ties, and intensified social competition (Gruebner et al., 2017). Epidemiological evidence, primarily derived from WEIRD (Western, educated, industrialized, rich, and democratic) populations, indicates that urban living is associated with an increased risk of depression and anxiety (Peen et al., 2010), as well as a dose–response relationship between urbanicity and schizophrenia (Gruebner et al., 2017). These risks are unevenly distributed due to structural inequalities, with socioeconomically disadvantaged and migrant populations experiencing disproportionate exposure through residential concentration in areas with inadequate infrastructure and limited protective resources (Christie-Mizell, 2022; Feldman and Steptoe, 2004; Rethy et al., 2021). Individual characteristics also modulate how urban life is experienced: women and gender minorities frequently report greater fear of crime and reduced perceptions of safety (Beebeejaun, 2017), while personality traits such as extraversion and emotional stability are linked to lower stress appraisals and more positive affective responses (Ebstrup et al., 2011; Şahin and Çetin, 2017). Taken together, these findings highlight the ambivalent impact of urban environments and how they shape mental health through a complex interplay of structural, social, and individual factors. In previous literature, urban environments are often considered in contrast to natural ones, which in turn have been shown to have positive effects on affective and cognitive outcomes (Bratman et al., 2012). In urban environments, natural elements like trees and plants are generally viewed as having a restorative/protective effect. Stress Reduction Theory (SRT; Ulrich, 1981) provides an account of the physiological and emotional effects humans experience upon exposure to nature. Over the course of evolution, natural environments have provided favorable conditions for human survival, thereby fostering an inherently positive perception of natural elements. However, restorative effects of nature are likely not solely attributable to hardwired, bottom-up responses shaped by evolution. Rather, an evolutionary-constructivist perspective posits that top-down cognitive processes, such as learned associations and individually acquired meanings related to nature, probably significantly moderate the extent to which individuals benefit from natural environments (Koivisto et al., 2022).

### 1.2. Neuroimaging Evidence of Environmental Impact

Neuro-Urbanism (Adli et al., 2017), an emergent domain within the wider research field of Environmental Neuroscience (Berman et al., 2019; Kühn and Gallinat, 2024), seeks to unravel how specific built and natural features of the environment modulate stress and restoration processes, with a focus on urban environments. By shaping affective responses, these mechanisms provide a crucial link between urban exposures and long-term physical as well as mental health outcomes. Leveraging neuroimaging tools such as electroencephalography (EEG) and magnetic resonance imaging (MRI) allows for investigating how urban exposures shape brain structure, function, and dynamics to provide empirical evidence guiding a health-promoting future urban development (Beyer et al., 2024; Grassini, 2024). Previous MRI studies provide evidence that urban upbringing is associated with heightened perigenual anterior cingulate cortex (pACC) activity and reduced pACC–amygdala connectivity, indicating impaired top-down regulation of affective stress (Lederbogen et al., 2011), and potentially reflecting a sustained state of hyper-vigilance (Kühn et al., 2021).

### 1.3. Previous EEG evidence

EEG studies corroborate these findings by documenting differential brain activity during exposure to natural versus built environments. Natural scenes are consistently rated as more relaxing, restorative, and fascinating than urban scenes (Elsadek et al., 2020; Hassan et al., 2018; Mahamane et al., 2020). In the EEG time domain, event-related potential (ERP) studies show that natural scenes elicit enhanced amplitudes of the early visual evoked (P1) component, potentially reflecting facilitated perceptual processes (Grassini et al., 2019). The P1 and N1 components, typically peaking within the first 200 milliseconds after stimulus onset at occipital electrode locations, have been widely replicated as being modulated by directed attention to visual stimuli in visuo-spatial attention paradigms (Luck and Kappenman, 2013). Both P1 and N1 components are sensitive to physical attributes of visual stimuli, including low-level visual features (e.g., luminance, color, contrast; Hillyard and Münte, 1984; Luck and Kappenman, 2013; Zhang and Luo, 2012). Together, the components are believed to represent top-down modulation of incoming sensory activity in the visual cortex (Luck and Kappenman, 2013). A functional dissociation between P1 and N1 is hypothesized, with the P1 component reflecting a sensory gain modulation of stimuli in attended locations (Hillyard et al., 1998), while the subsequent N1 component is assumed to reflect discriminatory processes directed to the information at the attended locations (Vogel and Luck, 2000). The P1 and N1 components have further been shown to be sensitive to emotional stimuli, with both components showing stronger deflections in response to negatively and positively charged stimuli relative to neutral stimuli (Hajcak et al., 2013). These findings, however, have been most robust for paradigms employing either emotional faces or very extreme emotional images.

Another ERP component sensitive to emotional images, but also specifically to urban scenes in Neuro-Urbanistic research, is P3. The P3 is a late positive component with a maximum over parietal sites peaking approximately 300–500 ms after stimulus onset. The P3 is considered sensitive to a stimulus’ motivational significance. In particular, P3 amplitudes are more pronounced for infrequent, task-relevant stimuli, consistent with the view that the component indexes the allocation of attentional resources (Polich, 2012). More pronounced amplitudes of the P3 component are interpreted as reflecting greater cognitive effort during context updating (Grassini et al., 2019; Kok, 1997; Luck and Kappenman, 2013; Rushby et al., 2005). Importantly, P3 amplitudes are also modulated by both positive and negative emotional valence, linking this later component to motivational mechanisms in addition to cognitive updating demands (Luck and Kappenman, 2013). Another emotionally modulated ERP component previously examined in the context of Neuro-Urbanism is the late positive potential (LPP). This component refers to a sustained positive deflection extending beyond the P3 time window in centroparietal regions. Although evidence suggests a general modulation by emotional stimuli (regardless of valence, in line with effects on previously presented components), some evidence points to an association between LPP amplitudes and subjective arousal (Cuthbert et al., 2000; Schupp et al., 2004; Weinberg and Hajcak, 2010; Hajcak et al., 2013) as well as with biologically relevant content, such as threat or erotic images (Hajcak et al., 2013; Schupp et al., 2004; Briggs and Martin, 2009). Regarding the LPP, findings in previous Neuro-Urbanistic research remain inconsistent: While Grassini et al. (2019) observed enhanced amplitudes in response to natural environments, Mahamane et al. (2020) reported elevated amplitudes occurring in response to built environments, yielding conflicting interpretations about emotional processing and sustained affective engagement.

### 1.4. Limitations of prior studies

Despite important new insights, studies in the field of Neuro-Urbanism often come with several limitations. They usually maximize the contrast between green and built settings by presenting extreme stimuli (e.g., bamboo forest vs. dense inner-city environments: Hassan et al., 2018; urban vs. natural stimuli: Grassini et al., 2019) rather than capturing the mixed and continuous nature of real urban environments. Cities typically present a complex mixture of built and natural features, further modulated by factors such as sky view, vehicle presence, and social density. In previous studies, sky views reduced perceived stress while enhancing fascination and openness (Asgarzadeh et al., 2014; Kühn et al., 2023), whereas vehicle presence increased arousal and diminished preference (Knöll et al., 2019). High social density has been associated with heightened arousal and negative affect, though these effects might be buffered by sky openness or opportunities for movement (Evans, 1979; Mavros et al., 2022). Additionally, other studies have shown that the openness/enclosure or physical built density of urban environments also influenced participants’ well-being and preferences (Shi et al., 2014; Trossman Haifler and Fisher-Gewirtzman, 2024). Beyond the presence or absence of natural elements, the spatial layout of urban scenes may independently shape affective responses. Prospect-Refuge Theory (Appleton, 1975) offers a theoretical basis for understanding these effects, proposing that humans derive feelings of safety from environments that balance prospect (i.e., the ability to survey one’s surroundings) with refuge (i.e., a sense of protective enclosure; Dosen and Ostwald, 2016). Applied to urban scenes, spatial openness may therefore undermine feelings of safety and comfort by maximizing prospect while minimizing refuge (Dosen and Ostwald, 2016). Within this framework, metric depth can be understood as an index of the prospect dimension. Scenes with greater spatial depth afford more visual access to the environment and thus a higher degree of prospect. Building on these findings, the present study employed previously tested real-world images of different urban scenes in Berlin, Germany (Sander et al., 2024) with varying proportions of vegetation, built structures, and sky, expecting that scenes with greater amounts of greenery and sky would be rated as less arousing and less stressful, more positive in valence and higher in beauty, fascination, and openness, while scenes with more greenery additionally would elicit higher ratings of hominess, feelings of control, and safety.

A further limitation of previous studies is the lack of systematic attention to low-level visual features (LLVFs) such as spatial frequency, color, texture, straight edge density (SED), or fractal patterns (e.g., Hagerhall et al., 2008). Beyond the semantic contents of an urban scene, these low-level visual properties shape rapid ‘gist’ extraction of a scene (Oliva, 2005), which might impact the scene appraisal through efficient feature prioritization (Groen et al., 2016) more than extensive inspection (Sander et al., 2024). Recent empirical work shows that measures such as hue, saturation, lightness, entropy, or sky/green pixel ratios predict perceived naturalness and restorativeness (Celikors and Wells, 2022; Ma et al., 2023; Samaan et al., 2024), though predictive strength varied across contexts. Beyond quantifying the composition of urban environments, the present study examined a set of LLVFs as predictors in an explorative fashion. We hypothesized that these features would shape scene appraisals in ways that are distinct from, and not fully explained by, their correlations with semantic scene categories.

Regarding neural responses to visually presented scenes, prior ERP research provides a basis for specific predictions. Given evidence that natural scenes facilitate early perceptual processing (Grassini et al., 2019), scenes with greater amounts of natural elements are expected to elicit larger P1 amplitudes, reflecting enhanced early sensory processing. Further, scenes with more urban elements were predicted to evoke more negative-going N1 amplitudes, indicating discriminatory processes, as well as enhanced P3 amplitudes, possibly reflecting greater cognitive effort and/or higher motivational salience (Grassini et al., 2019; Polich, 2012). Finally, because LPP amplitudes are linked to sustained affective engagement, we expected more restorative environments to evoke larger LPP responses (Grassini et al., 2019). However, mixed findings in prior work (Mahamane et al., 2020) limit how clearly such effects can be interpreted.

In summary, the present study investigated the following hypotheses:

**H1** We expected a higher proportion of natural elements (i.e., greenery and sky), as compared to man-made elements, in images to lead to more positive subjective ratings.

**H2** We expected a higher proportion of natural elements, as compared to man-made elements, to lead to more pronounced P1 amplitudes and less pronounced N1 amplitudes at occipital electrodes, as well as enhanced P3 amplitudes at parietal electrodes.

**H3** We expected LPP amplitudes at parietal electrodes to be modulated differently by natural elements, as compared to man-made elements.

**H4** We expected subjective responses to be modulated by the stimuli’s LLVFs, with features characteristic of natural elements having a similar effect to these elements’ direct presence.

**H5** Similarly, we expected P1, N1, P3 and LPP amplitudes to be modulated by the stimuli’s LLVFs, reflecting possible explanatory pathways of effects through early visual processing.

**H6** We exploratively investigated the connection between ERP components and subsequent subjective ratings.

## 2. Methods

The current EEG study investigates human brain responses to authentic urban scenes with varied proportions of vegetation and built structures. We used both semantic segmentation and low-level feature analysis and accounted for individual variation by including gender and personality dimensions, which have previously been shown to influence subjective ratings and physiological responses, as covariates in the analyses. Additionally, in a second step, we exploratively investigated the connection between subjective scene appraisal and early neural responses immediately following their perception.

### 2.1. Participants

For this study, 72 participants were recruited via TU Berlin’s participant recruitment portal. The sample consisted of students and citizens signing up to participate in research studies. Participation was reimbursed with either study credits or 12 EUR per hour. To qualify, participants had to be 18 years or older, fluent in English or German, and to have normal or corrected-to-normal vision with contact lenses. Four participants had to be excluded due to technical problems during data recording and two participants were excluded as they did not adhere to the instructions (e.g. excessively looking away from the screen due to eye strain). Finally, three participants were excluded due to poor EEG data quality (more than 20% of channels rejected in pre-processing or over 75% of epochs rejected). The final analysis was thus performed on a sample of 63 participants (age *M* = 29.16 years, *SD* = 7.53, range = 19–61 years; 29 male, 33 female, 1 non-binary). Ethical approval was obtained from the Ethics Board of the Institute of Psychology and Ergonomics at Technische Universität Berlin (ethics protocol BPN_GRA_230,415).

An a priori simulation-based power analysis was calculated using the approach suggested by Kumle et al. (2021). Simulations used the smallest effect sizes of interest based on effect sizes observed in a previous online study with a very similar experimental protocol (Sander et al., 2024). The results indicated a minimum power of >0.76 across the three fixed effects of interest, with a sample size of n = 60. The sample size recruited based on these power simulations should thus have been sufficiently powered to observe the hypothesized effects.

### 2.2. Design

The experiment used a within-person design in which the independent variables were images of different urban areas varying in urban density, operationalized as the proportion of image area covered by different object classes (i.e., buildings, greenery, sky, cars, persons). Participants subjectively rated each stimulus on nine scales. We used the Self Assessment Manikin (SAM) with a 9-point Likert scale to assess the three dependent variables dominance, arousal, and valence (Bradley and Lang, 1994). In addition, a set of six exploratory subjective measures assessing perceived stress, openness, safety, beauty, hominess, and fascination used 5-point Likert scales. We also used the Ten Item Personality Measure (TIPI) (Gosling et al., 2003) in the original version and its validated German translation (Muck et al., 2011) to measure the Big-5 personality traits extraversion and emotional stability, which were included in the analyses as control variables. For additional statistical control, the city size where participants spent the first 15 years of their life was recorded with a 5-point scale by number of inhabitants (< 5.000; 5.000 – 20.000; 20.000 – 100.000; 100.000 – 1 million; >1 million).

Participants’ brain activity was recorded throughout the experiment using 64-channel EEG. While we also recorded electrodermal activity (EDA), electrocardiography (ECG), and eye tracking, the current study focuses on the EEG data only.

### 2.3. Materials

The stimuli used in this study consisted of 56 street view images, which were a subset of the most extremely rated images from a set of 100 stimuli used in the previous study from our group (Sander et al., 2024). All images were of locations in Berlin, Germany, with varying levels of building height, greenery, and traffic. Images were taken as screenshots from high-resolution 360-degree street-level imagery (Apple Lookaround; Apple Inc., 2022).

### 2.4. Procedure

The experiment took place during weekdays between 09:00-17:00 from April to July, 2024. Participants were seated alone in a windowless, temperature-controlled room with stable lighting conditions. The experiment was presented on a 27” (diagonal) monitor with a 60Hz refresh rate at a resolution of 2560×1440p. The monitor was located on a desk approximately 60 cm away from participants, who sat facing a wall. Psychtoolbox (Brainard, 1997) for MATLAB (The Mathworks Inc., Version 2023b) was used to implement the experiment. During the experiment, participants were presented with different urban stimuli and had to subsequently rate them on the nine subjective rating scales (arousal, valence, dominance, stress, openness, safety, beauty, hominess, and fascination). Each of the 56 stimuli was repeated nine times over the time course of the experiment to be rated on each of the nine scales, resulting in 504 trials. All stimuli were initially randomly assigned to one of two blocks, and within each block, stimulus-scale combinations were randomized individually for each participant. Stimuli were thus presented across two blocks of 28 stimuli each, separated by a 2-5 minute self-paced break to prevent fatigue effects. Each experimental trial consisted of participants being presented with a word pair for 1000ms, priming them to the scale they would be presented with to ensure attention to relevant image features and ease of consecutive rating (arousal: excited – calm; valence: happy – unhappy; dominance: controlled – in control; stress: relaxed – stressful; openness: narrow – open; safety: unsafe – safe; beauty: ugly – beautiful; hominess: alienated – at home; fascination: boring – fascinating). Subsequently, a fixation cross appeared for 500 ms, followed by a stimulus for 3000 ms. After the stimulus disappeared, the rating scale was presented until participants logged their rating using the computer mouse. At the beginning and end of the experiment, there was a 3 min baseline recording in which participants were looking at a white screen with an instruction to keep eyes open and directed at the monitor. The average duration of the experimental task was 90 minutes per participant. For an overview of the experimental procedure, see Figure 1.

**Figure 1:**
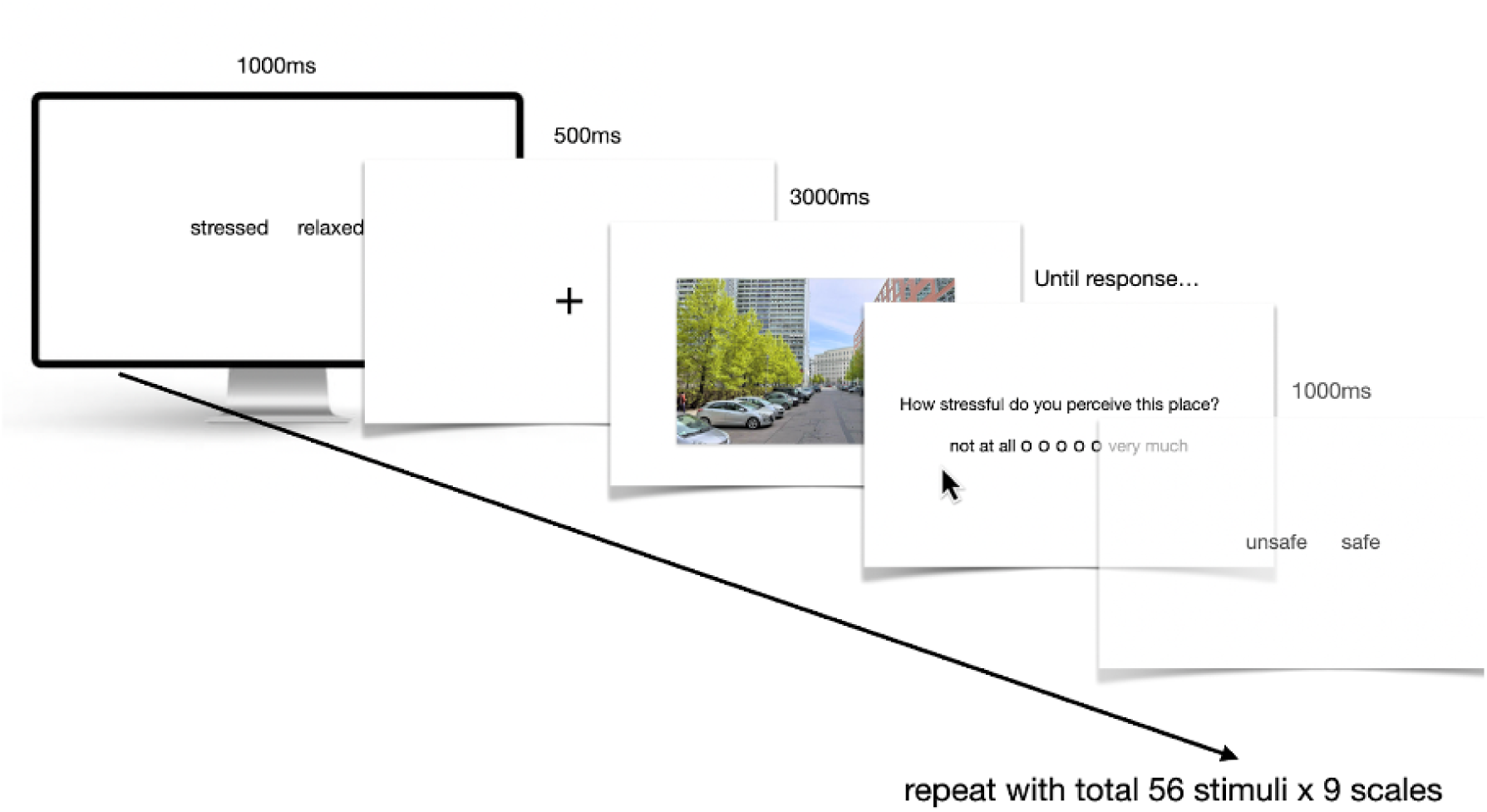
Schematic representation of the experimental trial structure. Each trial began with a word-pair prime indicating the subsequent rating dimension, followed by a fixation cross, an urban scene image, and a rating scale. The sequence was repeated for 56 stimuli across nine rating dimensions

### 2.5. Electroencephalography

Continuously recorded physiological data and event streams were synchronized and recorded using the Lab Streaming Layer (LSL; Kothe et al., 2025). EEG data were recorded continuously throughout the experiment from 64 actively amplified gel-based Ag/AgCl electrodes placed according to the extended 10% system (Chatrian et al., 1985) with FCz as reference and Fpz as ground. The data was sampled at 500 Hz with a 16-bit resolution using BrainAmp DC amplifiers from BrainProducts (BrainProducts GmbH, Gilching, Germany) with a 0.016 Hz high-pass filter during data acquisition. After fitting the cap, all electrodes were filled with conductive gel to ensure proper conductivity. Electrode impedances were kept below 10kΩ where possible.

#### 2.5.1. Preprocessing

Data preprocessing was performed using the ‘BeMoBIL Pipeline’ (Klug et al., 2022; https://github.com/BeMoBIL/bemobil-pipeline, v2.0.0, accessed 30.04.2024) implemented via the EEGLAB toolbox (Delorme and Makeig, 2004) for MATLAB (The Mathworks Inc., Version 2023b). After merging the two experimental blocks and removing non-experimental sections from the data, the EEG was filtered with a 125 Hz low-pass filter and downsampled to 250Hz. Subsequently, the data were subjected to the ZapLine-Plus function to remove spectral noise using the default automatic detection settings (Klug and Kloosterman, 2022). We then used the pipeline’s ‘bemobil_detect_bad_channels’ functions using default settings but lowering the maximum time during which a channel may be broken before rejection to 30% for a more aggressive detection approach due to overall good data quality. Rejected channels (*M* = 2.53, *SD* = 2.73) were subsequently interpolated using spherical spline interpolation, and the data were re-referenced to the average of all channels, including the reconstruction of the original reference channel FCz. After applying a zero-phase FIR high-pass filter (passband edge: 1.75Hz; –6dB cutoff: 1.5Hz; filter order: 1650), we used the BeMoBIL wrapper functions to perform Adaptive Mixture Independent Component Analysis (AMICA; Palmer et al., 2011) using the default parameters for AMICA, including 2,000 iterations and the sample rejection option. The high-pass filter and rejection were only applied to the data for AMICA computations. Equivalent dipole models were fitted for each of the resultant Independent Components (ICs) using the ‘dipfit’ toolbox of EEGLAB with default settings. The computed AMICA data and dipole models were copied back to the initial preprocessed dataset without any high-pass filter or rejected data. ICs were labeled using the ICLabel default popularity classifier, which assigns ICs one of seven labels (“brain,” “eye,” “muscle,” “heart,” “line noise,” “channel noise,” or “other”) (Pion-Tonachini et al., 2019). The final datasets were cleaned with ICA by removing ICs labeled as eye sources, and the data was lastly high-pass filtered with a zero-phase FIR filter (pass-band edge: 0.2Hz; –6dB cutoff: 0.1Hz; filter order: 4126).

### 2.6. Data Analysis

#### 2.6.1. Event-Related Potentials

The preprocessed EEG data were epoched using a time window of –200 to +1200 ms relative to stimulus onset. Baseline correction was applied using the interval from –200 to 0 ms. To reduce artifacts, noisy epochs were excluded if the amplitude exceeded ±100*µ*V in any channel, thereby removing trials contaminated by excessive transient signal fluctuations. To identify relevant ERP components, grand average waveforms were initially plotted and visually inspected. For early components (P1, N1), the grand-averaged signals were examined at parieto-occipital electrodes POz, Oz, O1, and O2, consistent with observed visual processing topographies (Colombari et al., 2024). For later components (P3, LPP), the centroparietal and parietal electrodes CPz, Pz, P3, and P4 were selected. This choice aligns with established findings that P3 and LPP components typically reach their peak amplitudes at centroparietal and parietal sites (Barry et al., 2020; Schupp et al., 2000; Hajcak et al., 2013). For the early components P1 and N1, grand average peak/trough latencies were extracted based on these visual inspections in the typical time ranges for each component (Luck and Kappenman, 2013), serving to define component-specific search time windows of ±20 ms around the identified maximum or minimum: The P1 component (peak identified at ∼105 ms) was analyzed within 85–125 ms, and the N1 component (peak identified at ∼152 ms) within 132–172 ms after stimulus onset. Component-specific peak detection was performed by automatically identifying the local maximum (P1) or minimum (N1) in these windows using a min/max search function. To improve robustness and to mitigate the influence of single-sample fluctuations, the final peak amplitude was calculated as the average across the identified peak and its two adjacent sample points (i.e., ±1 sample point or ±4 ms). For the later, more slowly varying components P3 and LPP, average amplitudes were extracted around the maxima identified from the grand average ERP waveform using previously published component time periods as guidance (Luck and Kappenman, 2013): P3 (maximum identified at ∼364 ms) within 339–389 ms, and the LPP (maximum identified at ∼535 ms) within 485–585 ms after stimulus onset, using a broader time-window for LPP to account for its more sustained nature.

The complete recording set-up and pre-processing of the EEG data has been registered on the ARTEM-IS platform (Styles et al., 2021)

#### 2.6.2. Quantification of Image contents

To obtain continuous quantification of the stimulus images’ contents, the images were first downsized to 1024×612px and then semantically segmented using the SegFormer-B5 model (Xie et al., 2021) pretrained on the ADE20K dataset (Zhou et al., 2017). Semantic segmentation is a machine-learning technique that assigns a class label to each pixel of an input image, effectively dividing the image into segments that represent various object classes. This allows for extracting the sums of pixels classified as being a specific object, in this case, buildings, greenery, cars, persons, and sky. Example outputs of the segmentation can be found in Figure 2. To evaluate the model’s segmentation performance, a subset of 12 images from the stimulus pool was manually labeled for the ADE20k object classes by two separate raters using Label Studio (Tkachenko et al., 2020-2025). Ground truth brushstrokes were exported and to compare the class masks with Segformer’s predictions, Intersect-over-Union (IoU) was calculated for the classes used as predictors in this study. This is the official evaluation metric used by the ADE20K benchmark (Zhou et al., 2017) and a multitude of other segmentation datasets (Wang et al., 2023), where pixel counts are pooled across all images before dividing, which provides more stable estimates in the presence of class imbalance across scenes. The model achieved a mean IoU (averaged across both raters) of 0.7 for the five classes. The lowest IoU was achieved for the “person” category with a value of 0.23. All other categories achieved an IoU ≥ 0.63. For the full results of both raters, please refer to the Supplemental Material. In the present analyses, the proportion of greenery was operationalized as a ratio of the sum of pixels classified as trees, plants, and grass relative to pixels classified as built structures (i.e., roads, sidewalks, towers, and buildings). This methodological decision was based on the observation of a pronounced negative correlation between the absolute pixel numbers of greenery and buildings, which would introduce multicollinearity into the linear models. Specifically, these two predictors represented mutually exclusive spatial features within a fixed two-dimensional image frame, resulting in a strong structural dependency. An analysis based on z-standardized values (N = 56) revealed a Pearson correlation coefficient of –0.81 (95% CI [-0.82, –0.80], *p* < .001), indicating a close inverse relationship. The predictors’ variance inflation factors (VIFs) in linear mixed models consistently exceeded the acceptable threshold of 5 when adding them separately (greenery: 5.15 – 5.63; buildings: 5.89 – 6.31), indicating problematic multicollinearity. To address this issue and obtain statistically interpretable estimates, we therefore computed a ratio of greenery relative to built structures. This relative metric reduces redundancy between predictors while retaining the ecological interpretability of the greenery variable within the given set of stimuli.

**Figure 2:**
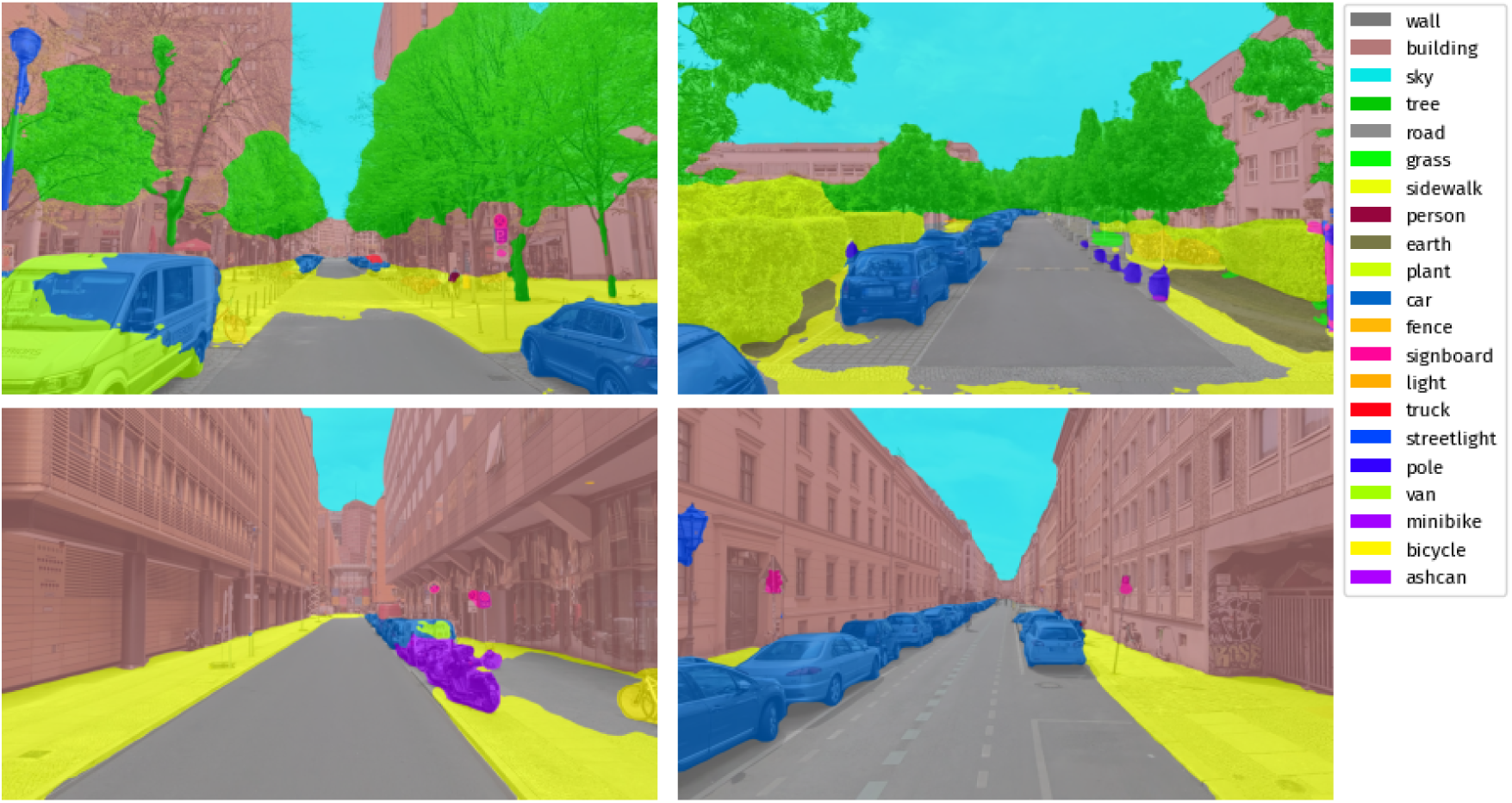
Example of four stimuli with varying levels of greenery and built density, sky, and traffic, shown with overlays of their respective semantic segmentation outputs. Top row, left: high greenery, high built density. Top row, right: high greenery, low built density. Bottom row, left: low greenery, high built density. Bottom row, right: low greenery, low built density.

#### 2.6.3. Low-Level Visual Features

Besides segmenting the images regarding the pixels occupied by built structures, greenery, cars, persons, and sky, we analyzed an exploratory set of LLVFs to provide insights into which aspects of the environment contribute to subjective experiences (Celikors and Wells, 2022; Samaan et al., 2024). A hierarchical model of scene processing suggests that early visual processing is primarily driven by LLVFs, which are basic image properties such as color information, contrast, and edge orientation, whereas later stages rely more heavily on object recognition and categorical representations (Groen et al., 2016). These LLVFs serve an adaptive function by facilitating rapid recognition and guiding navigational behavior in complex environments.

We extracted eight low-level visual features that have been shown to significantly influence natural scene perception in previous studies (Celikors and Wells, 2022; Ma et al., 2023; Samaan et al., 2024). Extraction of Straight (SED) and Non-Straight Edge Density (NSED), as well as entropy was adapted from Berman et al. (2014), with entropy serving as a measure of visual complexity. Color features were derived from the HSV color space. We computed the number of pixels in each image that fell into the green (60-180°) hue spectrum to quantify the images’ green color properties linked to natural and non-natural elements. To extract only pixels that can be reasonably considered perceptible as chromatic (i.e., colorful or perceivably different from gray/black/white), we set a perception cut-off according to the true color weight function (*W_H_* (*S, I*)) proposed by Vadivel et al. (2008), which models color perception based on pixels’ saturation and intensity values and includes perceptual tuning parameters based on human color perception. Pixels with true color weight values larger than 0.6 were considered reasonably perceivable for this analysis. We also included mean saturation values, as well as saturation’s standard deviation (SDsaturation). We additionally extracted images’ Fractal Dimension (FD), a non-integer value quantifying the complexity and self-similarity of a pattern. Stimuli’s mean spatial depth was estimated using the Depth Anything V2 model (Yang et al., 2024), specifically the outdoor-scene variant optimized for metric depth estimation. Together, this set of extracted features balances LLVF’s perceptual cues with higher-order spatial and textural information, laying the groundwork for linking continuous scene characteristics to neurophysiological evidence. For a distribution of all extracted LLVFs and their pairwise relationships, please refer to the Supplemental Material.

#### 2.6.4. Statistical Models

To examine the influence of visual stimulus features and individual differences on both subjective ratings and ERPs, a series of linear mixed models (LMMs) was computed using the lme4 package (Bates et al., 2015) in R (version 4.2.2; R Core Team, 2022). We focused the analyses on nine subjective outcome variables: arousal, valence, dominance, stress, safety, beauty, hominess, fascination, and openness, as well as amplitudes of four ERP components: P1, N1, P3, and LPP. Separate models were estimated for each dependent variable.

A total of 26 LMMs were fitted: two models for each dependent variable, one using predictors extracted from stimuli via semantic segmentation (greenery/built ratio, sky, car, person), and one using 8 measures of the stimuli’s LLVFs (saturation, SD of saturation, number of green pixels, entropy, SED, NSED, fractal dimension, mean of metric depth). To explore the connection between neural processes and subsequent subjective ratings, we also fitted nine linear models with the subjective ratings of interest as outcomes and mean ERP amplitudes as predictors. Each model additionally included individual-level control variables as fixed effects which were previously shown to influence appraisal of urban areas and/or physiological responses but were not of specific interest in the present study: gender, size of the city where the respondents spent the first 15 years of their life, extraversion, and emotional stability (Sander et al., 2024; Yuan et al., 2007; Hoffman and Polich, 1999). The gender variable was binarized to male and non-male due to a single participant reporting their self-identified gender as non-binary, which did not allow for separate statistical analysis. All continuous predictors were z-scaled prior to analysis due to differences in scales.

To account for the repeated measures design, we included random intercepts for both participant and stimulus (Judd et al., 2012). Where supported by model comparison, random slopes for key visual predictors were added to improve fit. For nested models, comparisons were conducted using likelihood-ratio tests (LRT, p < .05). The final model choice was guided by LRT results (for detailed results and final selected models see Supplemental Material). Statistical significance of fixed effects was assessed using the Satterthwaite approximation for degrees of freedom, as implemented in the lmerTest package (Kuznetsova et al., 2017).

## 3. Results

We examined the relationships between all semantic image contents, visual low-level features, and outcome variables (see Figure 3).

**Figure 3:**
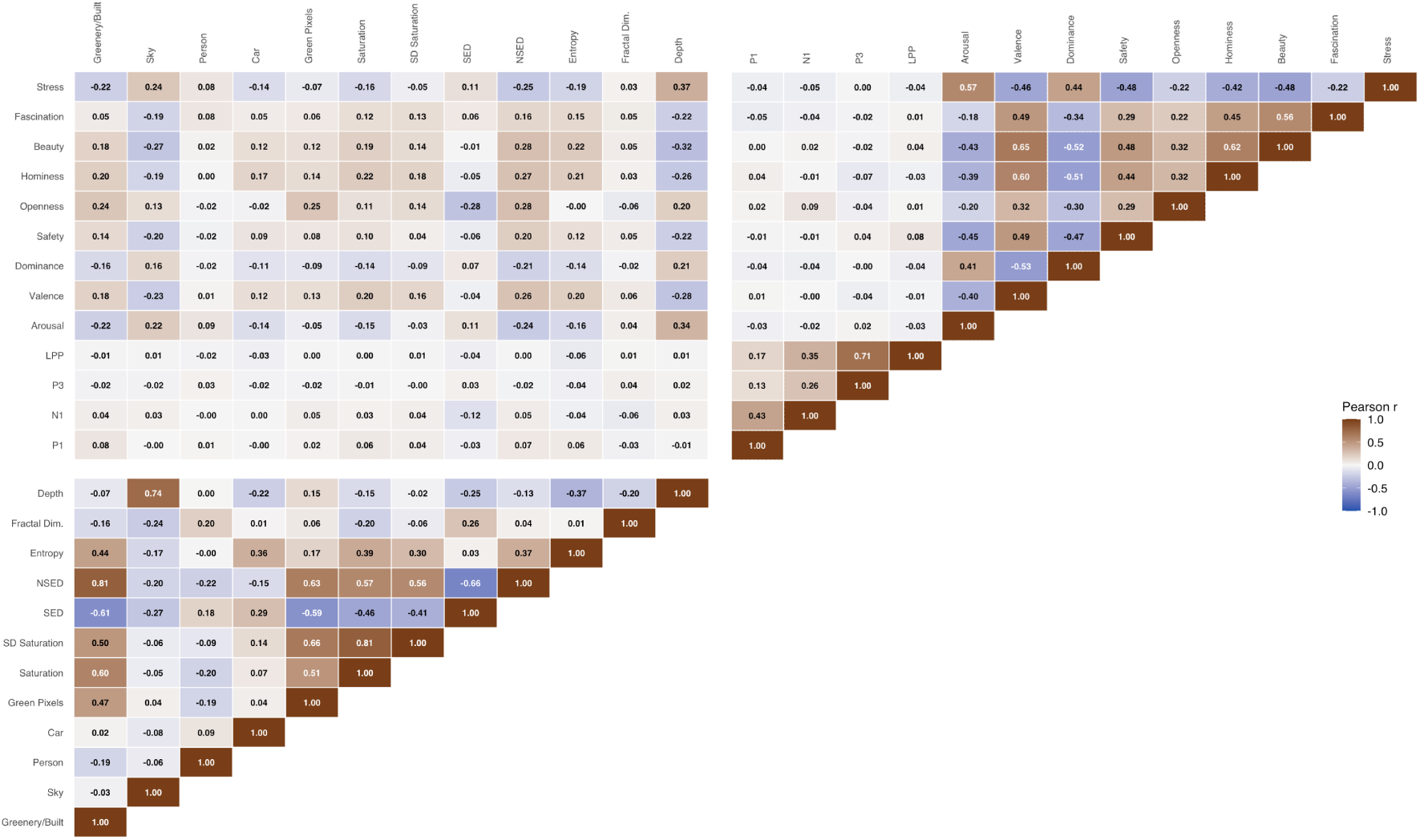
Correlation matrix showing Pearson correlations between all variables used in the study. The matrix includes semantic image features (greenery/built ratio, sky, person, car), low-level visual features (green pixels, saturation, SD of saturation, straight edge density [SED], non-straight edge density [NSED], entropy, fractal dimension, mean metric depth), ERP components (P1, N1, P3, LPP), and subjective ratings (arousal, valence, dominance, safety, openness, hominess, beauty, fascination, stress). Color coding ranges from blue (r = –1.0) to brown (r = 1.0). Strong correlations are highlighted in darker shades.

### 3.1. Subjective Measures

We report analyses for nine dimensions examining how visual and individual-level variables influence subjective evaluations. For an overview of rating distributions, see Figure 4.

**Figure 4:**
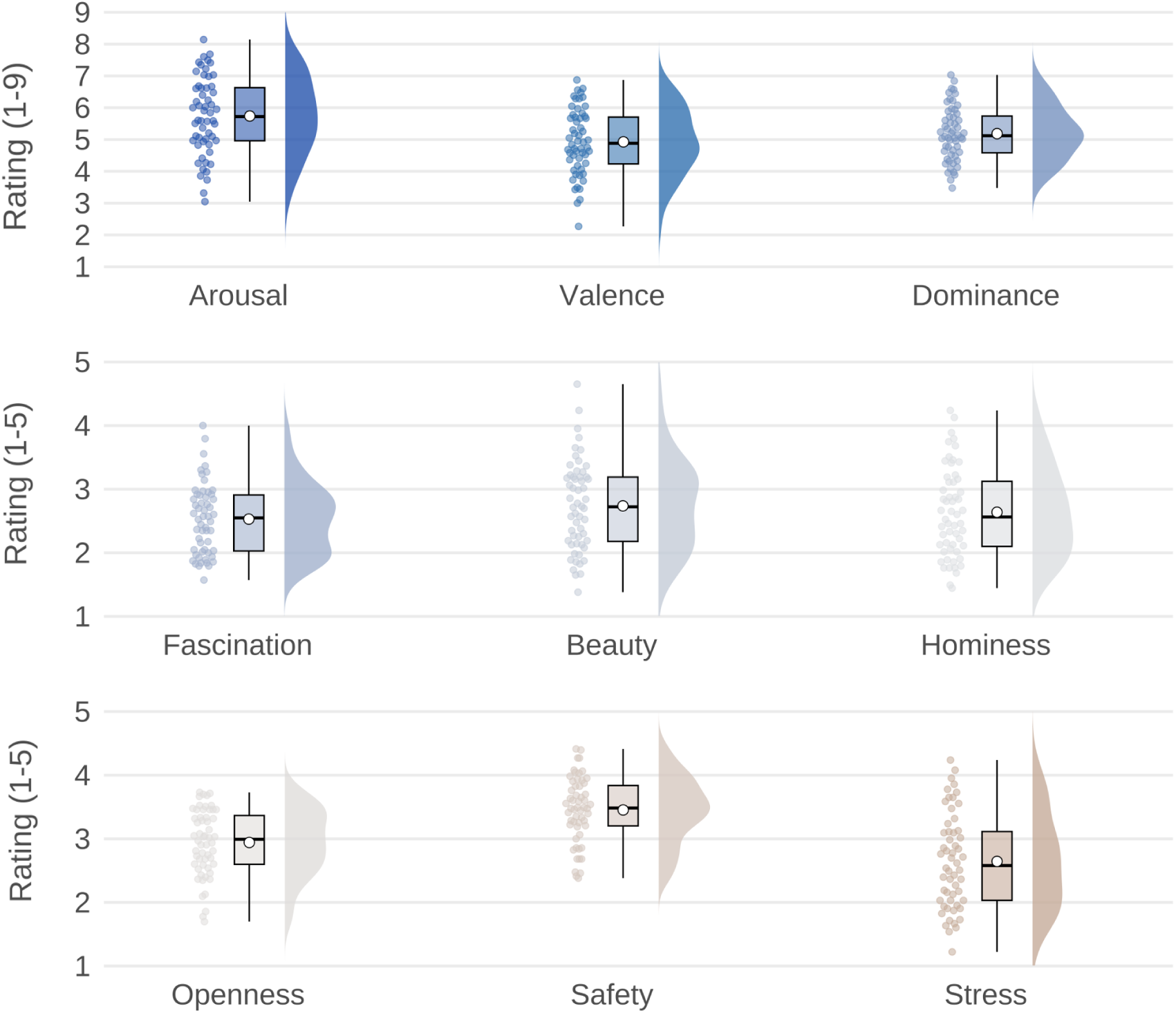
Raincloud plots of subjective ratings for SAM and Likert scales. Individual data points represent stimulus means aggregated across participants; boxplots show median (solid black line) and mean (white circle), while density distributions illustrate the variability across stimuli.

#### 3.1.1. Semantic Image Contents

The results of all LMMs using semantic segmentation categories as predictors and subjective ratings as outcomes are summarized in Tables 1 and 2. All *p*-values were adjusted for multiple hypothesis testing using the Benjamini-Hochberg procedure (FDR) (Benjamini and Hochberg, 1995).

**Table 1:** Results of LMMs predicting subjective ratings from semantic image contents(1/2).

| Predictors | Arousal |  |  |  | Valence |  |  |  | Dominance |  |  |  | Stress |  |  |  |
| --- | --- | --- | --- | --- | --- | --- | --- | --- | --- | --- | --- | --- | --- | --- | --- | --- |
|  | Est | SE | CI | *p*<sub>adj</sub> | Est | SE | CI | *p*<sub>adj</sub> | Est | SE | CI | *p*<sub>adj</sub> | Est | SE | CI | *p*<sub>adj</sub> |
| (Intercept) | 5.61 | 0.46 | 4.71-6.52 | <0.001 | 4.65 | 0.45 | 3.76-5.55 | <0.001 | 5.74 | 0.63 | 4.50-6.98 | <0.001 | 5.61 | 0.46 | 4.71-6.52 | <0.001 |
| greenery/built ratio | -0.41 | 0.15 | -0.69- -0.12 | **0.020** | 0.38 | 0.13 | 0.13-0.62 | **0.011** | -0.35 | 0.10 | -0.55- -0.14 | **0.005** | -0.41 | 0.15 | -0.69- -0.12 | **0.020** |
| sky | 0.43 | 0.14 | 0.16-0.70 | **0.014** | -0.44 | 0.12 | -0.67- -0.22 | **0.002** | 0.32 | 0.10 | 0.13-0.50 | **0.005** | 0.43 | 0.14 | 0.16-0.70 | **0.014** |
| car | -0.26 | 0.14 | -0.53-0.01 | 0.099 | 0.21 | 0.11 | -0.02-0.43 | 0.137 | -0.20 | 0.09 | -0.38- -0.02 | 0.081 | -0.26 | 0.14 | -0.53-0.01 | 0.099 |
| person | 0.16 | 0.14 | -0.12-0.44 | 0.304 | 0.06 | 0.12 | -0.17-0.28 | 0.701 | -0.07 | 0.09 | -0.25-0.12 | 0.609 | 0.16 | 0.14 | -0.12-0.44 | 0.304 |
| Gender [Non-male] | 0.15 | 0.17 | -0.17-0.48 | 0.358 | -0.30 | 0.17 | -0.63- -0.02 | 0.137 | 0.12 | 0.23 | -0.34-0.58 | 0.696 | 0.15 | 0.17 | -0.17-0.48 | 0.358 |
| City size first 15 years | -0.15 | 0.06 | -0.26- -0.04 | **0.020** | 0.06 | 0.06 | -0.05-0.17 | 0.381 | -0.09 | 0.08 | -0.24-0.07 | 0.410 | -0.15 | 0.06 | -0.26- -0.04 | **0.020** |
| Extraversion | -0.04 | 0.03 | -0.09-0.02 | 0.274 | 0.04 | 0.03 | -0.01-0.10 | 0.211 | -0.07 | 0.04 | -0.15-0.01 | 0.182 | -0.04 | 0.03 | -0.09-0.02 | 0.274 |
| Emotional Stability | -0.06 | 0.03 | -0.12- -0.01 | 0.064 | -0.00 | 0.03 | -0.06-0.06 | 0.992 | -0.01 | 0.04 | -0.09-0.07 | 0.819 | -0.06 | 0.03 | -0.12- -0.01 | 0.064 |
| Observations | 3528 |  |  |  | 3528 |  |  |  | 3528 |  |  |  | 3528 |  |  |  |
| Marginal R<sup>2</sup> / Conditional R<sup>2</sup> | 0.128 / 0.459 |  |  |  | 0.104 / 0.395 |  |  |  | 0.072 / 0.384 |  |  |  | 0.128 / 0.459 |  |  |  |

**Table 2:**
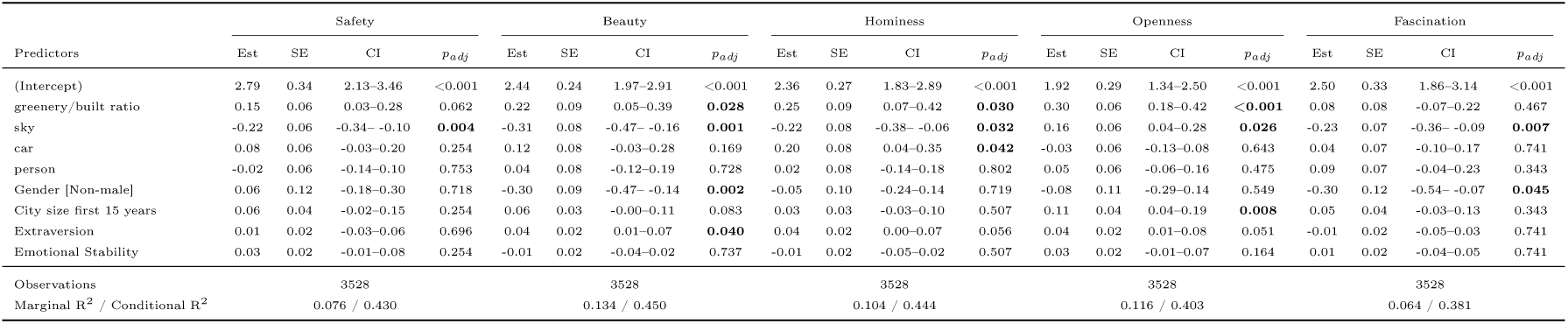
Results of LMMs predicting subjective ratings from semantic image contents(2/2)

The model for the *arousal ratings* showed that higher greenery/built ratios led to stimuli being rated as significantly more calm (*β* = –0.41, *p* = .02). Conversely, the amount of sky area in images had the opposite effect, with more sky area leading to higher arousal ratings (*β* = 0.43, *p* = .014). All other predictors did not significantly influence arousal ratings (all *p* ≥ .099).

Similar effects were observed for *valence ratings*. Greenery/built ratios exhibited a positive effect (*β* = 0.38, *p* = .011), suggesting that images containing more natural and less artificial elements were associated with more positive emotional evaluations. Sky area had a negative effect on valence ratings (*β* = –0.44, *p* = .002). None of the other predictors demonstrated effects on valence ratings (all *p* ≥ .137)

The *dominance ratings* assessed participants’ feeling of being in control vs. feeling controlled. The selected final model again showed that greenery/built ratios had a significant negative effect on perceived dominance (*β* = –0.35, *p* = .005), indicating that scenes with a higher proportion of vegetation compared to buildings and roads elicited stronger feelings of being in control. Sky area had a positive effect on perceived dominance ratings (*β* = 0.32, *p* = .005), reflecting higher feelings of being controlled. No other predictor showed effects on dominance ratings (all *p* ≥ .081.).

*Safety ratings* were significantly influenced only by the amount of sky in the images, with larger sky areas being perceived as less safe (*β* = –0.22, *p* = .004). No other predictor revealed an impact on safety ratings (all *p* ≥ .062).

The model for *beauty ratings* showed a significant effect of greenery/built ratio with images with more greenery compared to built areas being rated as more beautiful (*β* = 0.22, *p* = .028). On the other hand, larger sky areas made the images appear less beautiful (*β* = –0.31, *p* = .001). All other predictors did not significantly influence beauty ratings (all *p* ≥ .083).

As for *hominess ratings*, significant effects were found for greenery/built ratios with higher ratios positively influencing hominess ratings (*β* = 0.25, *p* = .030). Conversely, sky area had a negative effect on perceived hominess (*β* = –0.22, *p* = .032). Additionally, larger amounts of image areas covered by cars led to participants perceiving the scenes as homier (*β* = 0.20, *p* = .042). All other predictors did not significantly influence hominess ratings (all *p* ≥ .056).

*Openness ratings* showed significant positive effects of greenery/built ratios (*β* = 0.30, p < .001), as well as sky area (*β* = 0.16, *p* = .026). All other predictors did not significantly influence openness ratings (all *p* ≥ .051).

Of the semantic image contents, *fascination ratings* were only significantly affected by sky area, with more visible sky leading to lower fascination ratings (*β* = –0.23, *p* =0.007). All other predictors did not significantly influence fascination ratings (all *p* ≥ .343).

The final LMM analyzing the influence of visual features on *stress ratings* also showed a significant negative effect of greenery/built ratios (*β* = –0.41, *p* = .020), showing images with proportionally more greenery compared to buildings and roads being rated as less stressful. Sky area had the opposite effect, suggesting more visible sky area to be perceived as more stressful (*β* = 0.43, *p* = .014). All other predictors had no significant effects (all *p* ≥ .064).

#### 3.1.2. Lower-level visual Features

Tables 3 and 4 show the results of all models using LLVFs as predictors and subjective ratings as outcomes. All *p*-values were adjusted for multiple hypothesis testing using FDR.

**Table 3:** Results of LMMs predicting subjective ratings from LLVFs of images (1/2). *Note: CI are 95% confidence intervals; p-values were adjusted using FDR method; all predictors were z-scaled*.

| Predictors | Arousal |  |  |  | Valence |  |  |  | Dominance |  |  |  | Stress |  |  |  |
| --- | --- | --- | --- | --- | --- | --- | --- | --- | --- | --- | --- | --- | --- | --- | --- | --- |
| | Est | SE | CI | $p_{adj}$ | Est | SE | CI | $p_{adj}$ | Est | SE | CI | $p_{adj}$ | Est | SE | CI | $p_{adj}$ |
| (Intercept) | 5.59 | 0.46 | 4.70-6.49 | <0.001 | 4.73 | 0.46 | 3.83-5.62 | <0.001 | 5.66 | 0.63 | 4.43-6.90 | <0.001 | 3.50 | 0.25 | 3.02-3.98 | <0.001 |
| Green Hue | -0.09 | 0.19 | -0.47-0.28 | 0.749 | 0.14 | 0.16 | -0.17-0.45 | 0.549 | -0.08 | 0.14 | -0.35-0.19 | 0.717 | -0.08 | 0.12 | -0.31-0.14 | 0.717 |
| Saturation | -0.23 | 0.23 | -0.69-0.23 | 0.425 | 0.23 | 0.19 | -0.14-0.61 | 0.425 | -0.15 | 0.17 | -0.48-0.18 | 0.603 | -0.13 | 0.14 | -0.41-0.14 | 0.603 |
| SD of Saturation | 0.51 | 0.24 | 0.04-0.97 | 0.084 | -0.15 | 0.19 | -0.53-0.23 | 0.568 | 0.19 | 0.17 | -0.14-0.52 | 0.593 | 0.29 | 0.14 | 0.01-0.57 | 0.593 |
| Straight Edge Density | 0.06 | 0.21 | -0.35-0.48 | 0.765 | 0.39 | 0.17 | 0.05-0.72 | 0.094 | -0.14 | 0.15 | -0.43-0.16 | 0.603 | 0.06 | 0.13 | -0.19-0.31 | 0.603 |
| Non-Straight Edge Density | -0.47 | 0.21 | -0.89- -0.05 | 0.084 | 0.61 | 0.17 | 0.27-0.95 | **0.004** | -0.44 | 0.15 | -0.74- -0.14 | **0.027** | -0.24 | 0.13 | -0.49-0.01 | 0.265 |
| Entropy | 0.05 | 0.15 | -0.24-0.34 | 0.765 | -0.06 | 0.12 | -0.29-0.18 | 0.718 | 0.04 | 0.11 | -0.17-0.25 | 0.776 | 0.00 | 0.09 | -0.17-0.18 | 0.776 |
| Fractal Dimension | 0.21 | 0.14 | -0.06-0.47 | 0.255 | -0.05 | 0.11 | -0.27-0.17 | 0.718 | 0.07 | 0.10 | -0.12-0.26 | 0.699 | 0.11 | 0.08 | -0.05-0.27 | 0.699 |
| Mean metric depth | 0.70 | 0.15 | 0.41-1.00 | <**0.001** | -0.43 | 0.12 | -0.67- -0.19 | **0.004** | 0.38 | 0.11 | 0.17-0.60 | **0.007** | 0.46 | 0.09 | 0.28-0.63 | **0.007** |
| Gender binary [Non-male] | 0.17 | 0.17 | -0.15-0.49 | 0.425 | -0.29 | 0.17 | -0.62-0.04 | 0.230 | 0.11 | 0.23 | -0.35-0.57 | 0.768 | 0.06 | 0.09 | -0.12-0.23 | 0.768 |
| City size first 15 years | -0.15 | 0.06 | -0.26- -0.04 | **0.042** | 0.06 | 0.06 | -0.05-0.17 | 0.521 | -0.10 | 0.08 | -0.26-0.05 | 0.511 | -0.07 | 0.03 | -0.13- -0.01 | 0.511 |
| Extraversion | -0.04 | 0.03 | -0.09-0.02 | 0.339 | 0.04 | 0.03 | -0.02-0.10 | 0.409 | -0.07 | 0.04 | -0.15-0.01 | 0.331 | -0.04 | 0.02 | -0.07- -0.01 | 0.331 |
| Emotional Stability | -0.06 | 0.03 | -0.12- -0.01 | 0.084 | -0.00 | 0.03 | -0.06-0.06 | 0.915 | 0.00 | 0.04 | -0.08-0.08 | 0.964 | -0.03 | 0.02 | -0.06-0.00 | 0.964 |
| Observations | 3528 |  |  |  | 3528 |  |  |  | 3528 |  |  |  | 3528 |  |  |  |
| Marginal $R^2$ / Conditional $R^2$ | 0.201 / 0.473 | | | | 0.154 / 0.377 | | | | 0.090 / 0.380 | | | | 0.221 / 0.482 | | | |

**Table 4:** Results of LMMs predicting subjective ratings from LLVFs of images (2/2). *Note: CI are 95% confidence intervals; p-values were adjusted using FDR method; all predictors were z-scaled*.

| Predictors | Safety |  |  |  | Beauty |  |  |  | Hominess |  |  |  | Openness |  |  |  | Fascination |  |  |  |
| --- | --- | --- | --- | --- | --- | --- | --- | --- | --- | --- | --- | --- | --- | --- | --- | --- | --- | --- | --- | --- |
| | Est | SE | CI | $p_{adj}$ | Est | SE | CI | $p_{adj}$ | Est | SE | CI | $p_{adj}$ | Est | SE | CI | $p_{adj}$ | Est | SE | CI | $p_{adj}$ |
| (Intercept) | 2.81 | 0.34 | 2.15-3.47 | <0.001 | 2.44 | 0.24 | 1.98-2.91 | <0.001 | 2.35 | 0.27 | 1.83-2.88 | <0.001 | 1.94 | 0.30 | 1.36-2.52 | <0.001 | 2.50 | 0.32 | 1.87-3.14 | <0.001 |
| Green Hue | 0.10 | 0.09 | -0.08-0.28 | 0.528 | 0.12 | 0.11 | -0.09-0.33 | 0.345 | 0.08 | 0.12 | -0.16-0.32 | 0.701 | 0.07 | 0.06 | -0.05-0.19 | 0.451 | 0.04 | 0.09 | -0.14-0.23 | 0.810 |
| Saturation | 0.10 | 0.11 | -0.12-0.32 | 0.528 | 0.14 | 0.13 | -0.12-0.39 | 0.345 | 0.13 | 0.15 | -0.16-0.43 | 0.626 | -0.07 | 0.07 | -0.22-0.07 | 0.474 | 0.00 | 0.11 | -0.22-0.23 | 0.994 |
| SD of Saturation | -0.22 | 0.11 | -0.44-0.00 | 0.188 | -0.15 | 0.13 | -0.41-0.11 | 0.345 | -0.07 | 0.15 | -0.36-0.23 | 0.745 | -0.03 | 0.08 | -0.18-0.11 | 0.711 | 0.08 | 0.12 | -0.15-0.31 | 0.701 |
| Straight Edge Density | 0.10 | 0.10 | -0.10-0.29 | 0.528 | 0.34 | 0.12 | 0.11-0.57 | **0.015** | 0.22 | 0.13 | -0.05-0.48 | 0.289 | 0.00 | 0.07 | -0.13-0.13 | 0.969 | 0.34 | 0.10 | 0.14-0.55 | **0.008** |
| Non-Straight Edge Density | 0.28 | 0.10 | 0.08-0.48 | **0.035** | 0.47 | 0.12 | 0.24-0.71 | **0.002** | 0.35 | 0.14 | 0.09-0.62 | 0.058 | 0.37 | 0.07 | 0.24-0.51 | <**0.001** | 0.35 | 0.10 | 0.15-0.56 | **0.008** |
| Entropy | -0.05 | 0.07 | -0.19-0.09 | 0.591 | -0.05 | 0.08 | -0.21-0.11 | 0.570 | -0.00 | 0.09 | -0.19-0.18 | 0.961 | -0.03 | 0.05 | -0.12-0.06 | 0.609 | -0.05 | 0.07 | -0.20-0.09 | 0.701 |
| Fractal Dimension | -0.02 | 0.07 | -0.15-0.11 | 0.770 | -0.08 | 0.08 | -0.23-0.06 | 0.345 | -0.05 | 0.09 | -0.23-0.12 | 0.701 | -0.06 | 0.04 | -0.14-0.03 | 0.382 | -0.07 | 0.07 | -0.20-0.06 | 0.576 |
| Mean metric depth | -0.21 | 0.07 | -0.35- -0.07 | **0.027** | -0.29 | 0.08 | -0.46- -0.13 | **0.003** | -0.23 | 0.09 | -0.42- -0.05 | 0.058 | 0.23 | 0.05 | 0.13-0.34 | <**0.001** | -0.18 | 0.07 | -0.32- -0.04 | **0.039** |
| Gender binary [Non-male] | 0.04 | 0.12 | -0.21-0.28 | 0.770 | -0.29 | 0.09 | -0.46- -0.13 | **0.003** | -0.04 | 0.10 | -0.23-0.15 | 0.745 | -0.09 | 0.11 | -0.30-0.13 | 0.563 | -0.30 | 0.12 | -0.54- -0.07 | **0.039** |
| City size first 15 years | 0.06 | 0.04 | -0.02-0.14 | 0.392 | 0.06 | 0.03 | -0.00-0.11 | 0.113 | 0.03 | 0.03 | -0.04-0.09 | 0.626 | 0.11 | 0.04 | 0.04-0.19 | **0.011** | 0.05 | 0.04 | -0.03-0.13 | 0.417 |
| Extraversion | 0.02 | 0.02 | -0.03-0.06 | 0.591 | 0.04 | 0.02 | 0.01-0.07 | **0.044** | 0.04 | 0.02 | 0.01-0.08 | 0.058 | 0.05 | 0.02 | 0.01-0.08 | 0.058 | -0.01 | 0.02 | -0.05-0.03 | 0.810 |
| Emotional Stability | 0.03 | 0.02 | -0.01-0.07 | 0.392 | -0.01 | 0.02 | -0.04-0.02 | 0.710 | -0.02 | 0.02 | -0.05-0.02 | 0.626 | 0.03 | 0.02 | -0.01-0.07 | 0.308 | 0.01 | 0.02 | -0.04-0.05 | 0.810 |
| Observations | 3528 |  |  |  | 3528 |  |  |  | 3528 |  |  |  | 3528 |  |  |  | 3528 |  |  |  |
| Marginal $R^2$ / Conditional $R^2$ | 0.102 / 0.437 | | | | 0.212 / 0.438 | | | | 0.143 / 0.428 | | | | 0.173 / 0.389 | | | | 0.118 / 0.361 | | | |

*Arousal ratings* showed significant effects of metric depth of the urban scenes. A higher mean depth of a scene led to significantly higher arousal ratings (*β* = 0.70, p < .001). All other LLVFs had no significant effect on subjective arousal (all *p* ≥ .084).

The model for *valence ratings* showed that participants reported themselves significantly happier in scenes with higher non-straight edge density (*β* = 0.61, *p* = .004), indicating a positive impact of more unstructured (natural) edges in the scenes. Additionally, metric depth once again had a significant effect; higher mean depth significantly decreased valence ratings (*β* = –0.43, *p* = .004). None of the other LLVFs had a significant effect on valence ratings (all *p* ≥ .094).

Similar to valence ratings, *dominance ratings* were also significantly affected by scenes’ mean metric depth and non-straight edge density. Higher non-straight edge density was associated with participants reporting feeling more in control (*β* = – 0.44, *p* = .027), while higher depths were associated with higher feelings of being controlled (*β* = 0.38, *p* = .007). Other LLVFs had no effects on dominance ratings (all *p* ≥ .593)

*Safety ratings* were significantly affected by non-straight edge density and mean metric depth. More non-straight edges led to higher perceived safety (*β* = 0.28, *p* = .035), while higher mean metric depth led to lower safety ratings (*β* = –0.21, *p* = .027). No other LLVF showed significant effects on scenes’ safety ratings (all *p* ≥ .188).

For *beauty ratings*, the model showed a significant positive effect of both straight edge density (*β* = 0.34, *p* = .015) and non-straight edge density (*β* = 0.47, *p* = .002). Additionally, beauty ratings were significantly influenced by mean metric depth, with higher depths in images leading to lower beauty scores (*β* = –0.29, *p* = .003). No other LLVF showed significant effects on perceived beauty (all *p* ≥ .345).

None of the LLVFs had significant effects on *hominess ratings* (all *p* ≥ .058).

The model for perceived *openness ratings* revealed a significant positive effect of non-straight edge density (*β* = 0.37, p < .001) as well as mean metric depth (*β* = 0.23, p < .001). Other LLVFs had no significant effect (all *p* ≥ .382).

*Fascination ratings*, similar to beauty ratings, were positively affected by both straight- (*β* = 0.34, *p* = .008) and non-straight edge density (*β* = 0.35, *p* = .008). The model also showed a significant negative effect of mean metric depth (*β* = –0.18, *p* = .039). Other LLVFs, again, had no significant effect (all *p* ≥ .576).

Lastly, *stress ratings* were once more only significantly impacted by mean metric depth: higher depths in images led to participants rating them as more stressful (*β* = 0.46, *p* = .007). All other LLVFs had no significant effects on stress ratings (all *p* ≥ .265).

In summary, of all eight extracted LLVFs, only edge density measures and mean metric depth significantly affected subjective scene perception.

### 3.2. EEG

Grand average potentials across all images and participants revealed clear and pronounced ERP components with a visual evoked P1-N1 complex over occipital sites and a P3 component followed by an LPP complex over centroparietal sites (See figure 5). To evaluate the influence of visual predictors on neural responses to environmental stimuli, these four ERP components were analyzed. For each component, hierarchical LMMs were fitted in the same way as was done for the subjective outcome measures. All *p*-values for models with shared predictor sets were adjusted for multiple hypothesis testing using FDR.

**Figure 5:**
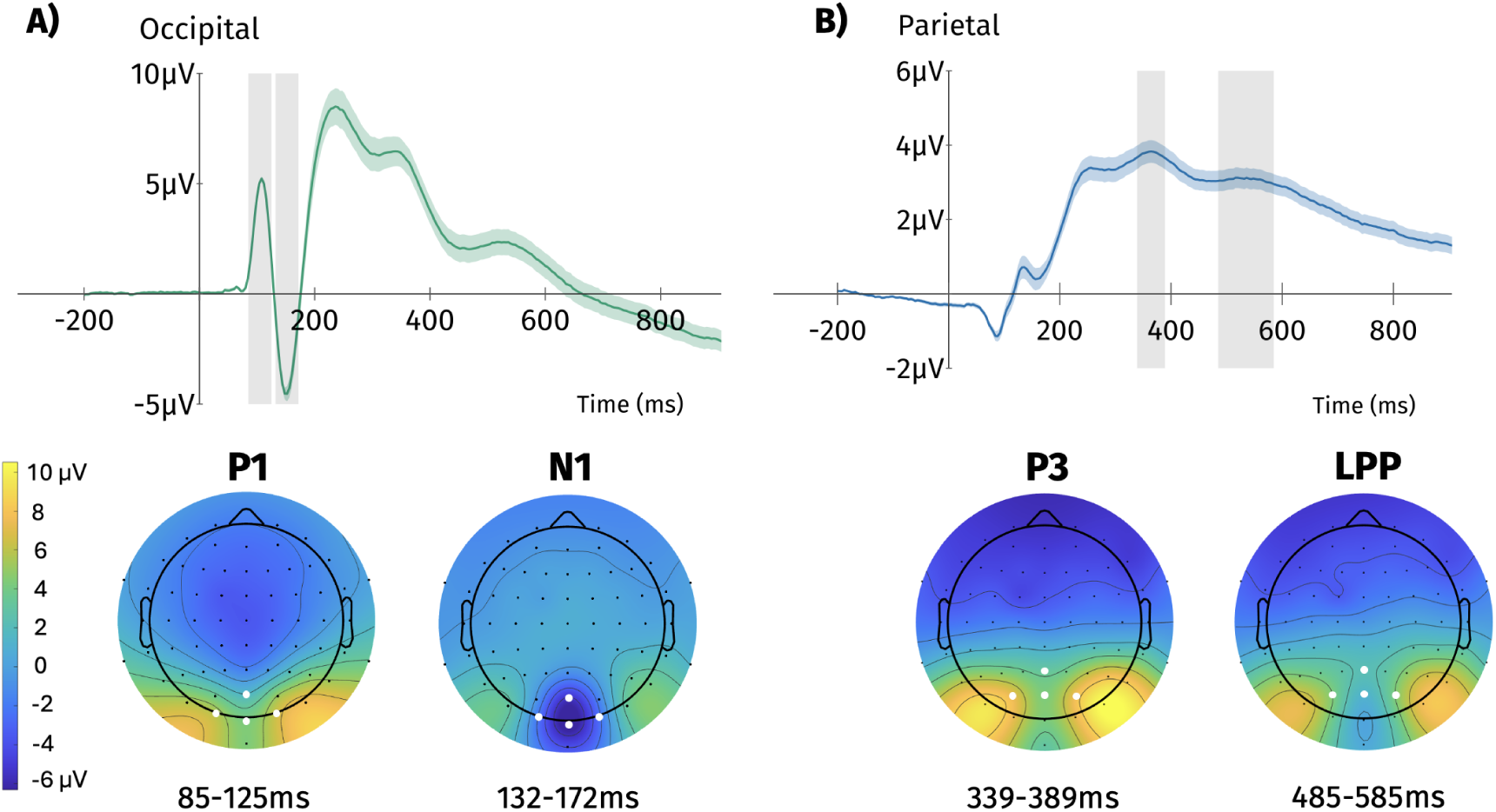
Grand Average ERP waveforms (averaged across all participants and stimuli) at onset of urban scene stimuli for the (A) occipital and the (B) parietal cluster. Color-shaded areas represent ± 1 standard error of the mean (SEM). (C) Bottom of Figure shows topographical plots of averaged amplitude activity (in *µ*V) during the time windows (indicated by the grey-shaded time periods) of the four extracted ERP components. *Note differences in scale of y-axes!*

#### 3.2.1. Semantic Image Contents

The LMM showed a significant effect of greenery/built ratio on occipital *P1-amplitudes* (*β* = 0.54, *p* = .001). Higher proportions of natural compared to artificial elements were associated with more pronounced P1-amplitudes. All other stimuli characteristics had no significant effects on P1 amplitudes (all *p* ≥ .288).

The subsequent *N1 component* did not reveal any significant effects (all *p* ≥ .433) regarding the semantic image contents.

For the *P3 and LPP components* at the centroparietal electrode cluster, none of the image contents reached statistical significance (all *p* ≥ .752). Full results can be found in Table 5.

**Table 5:** Results of LMMs predicting ERP component amplitudes from semantic image contents. *Note: CI are 95% confidence intervals; p-values were adjusted using FDR method; all predictors were z-scaled*.

| Predictors | mean P1 |  |  |  | mean N1 |  |  |  | mean P3 |  |  |  | mean LPP |  |  |  |
| --- | --- | --- | --- | --- | --- | --- | --- | --- | --- | --- | --- | --- | --- | --- | --- | --- |
|  | Est | SE | CI | *p*<sub>adj</sub> | Est | SE | CI | *p*<sub>adj</sub> | Est | SE | CI | *p*<sub>adj</sub> | Est | SE | CI | *p*<sub>adj</sub> |
| (Intercept) | 13.95 | 3.16 | 7.75–20.16 | <0.001 | -7.78 | 4.42 | -16.44–0.89 | 0.376 | 4.35 | 1.70 | 1.03–7.68 | 0.115 | 1.76 | 1.54 | -1.26–4.79 | 0.762 |
| greenery/built ratio | 0.54 | 0.14 | 0.26–0.81 | **0.001** | 0.37 | 0.26 | -0.15–0.89 | 0.433 | -0.06 | 0.09 | -0.24–0.12 | 0.752 | -0.04 | 0.10 | -0.23–0.15 | 0.873 |
| sky | 0.01 | 0.12 | -0.23–0.26 | 0.905 | 0.28 | 0.26 | -0.22–0.78 | 0.461 | -0.08 | 0.09 | -0.26–0.10 | 0.752 | 0.02 | 0.10 | -0.17–0.20 | 0.878 |
| car | -0.05 | 0.12 | -0.29–0.20 | 0.806 | 0.04 | 0.26 | -0.46–0.54 | 0.882 | -0.07 | 0.09 | -0.25–0.11 | 0.752 | -0.10 | 0.10 | -0.29–0.09 | 0.762 |
| person | 0.18 | 0.13 | -0.07–0.43 | 0.288 | 0.07 | 0.26 | -0.44–0.58 | 0.880 | 0.07 | 0.09 | -0.11–0.26 | 0.752 | -0.05 | 0.10 | -0.24–0.14 | 0.873 |
| Gender binary [Non-male] | 0.54 | 1.18 | -1.78–2.86 | 0.806 | -4.79 | 1.65 | -8.03–-1.55 | **0.047** | -0.90 | 0.63 | -2.14–0.34 | 0.724 | -0.56 | 0.58 | -1.69–0.57 | 0.762 |
| City size first 15 years | -0.57 | 0.40 | -1.36–0.22 | 0.288 | 0.74 | 0.56 | -0.36–1.84 | 0.433 | 0.00 | 0.22 | -0.42–0.42 | 0.991 | 0.25 | 0.20 | -0.14–0.63 | 0.762 |
| Extraversion | 0.10 | 0.21 | -0.31–0.51 | 0.806 | 0.22 | 0.29 | -0.36–0.79 | 0.590 | -0.02 | 0.11 | -0.24–0.20 | 0.991 | 0.07 | 0.10 | -0.13–0.27 | 0.873 |
| Emotional Stability | -0.37 | 0.21 | -0.79–0.04 | 0.256 | -0.31 | 0.30 | -0.89–0.28 | 0.461 | 0.00 | 0.11 | -0.22–0.23 | 0.991 | 0.02 | 0.10 | -0.19–0.22 | 0.878 |
| Observations | 3528 |  |  |  | 3528 |  |  |  | 3493 |  |  |  | 3493 |  |  |  |
| Marginal R<sup>2</sup> / Conditional R<sup>2</sup> | 0.053 / 0.660 |  |  |  | 0.116 / 0.792 |  |  |  | 0.020 / 0.580 |  |  |  | 0.026 / 0.543 |  |  |  |

#### 3.2.2. Lower-level visual Features

The models with LLVFs as predictors revealed only one significant effect: higher straight edge density led to more pronounced *N1 amplitudes* (*β* = –1.22, *p* = .024). None of the other LLVFs had a significant effect on any of the other three ERP components (all *p* ≥ 0.075). For full results, see Table 6.

**Table 6:**
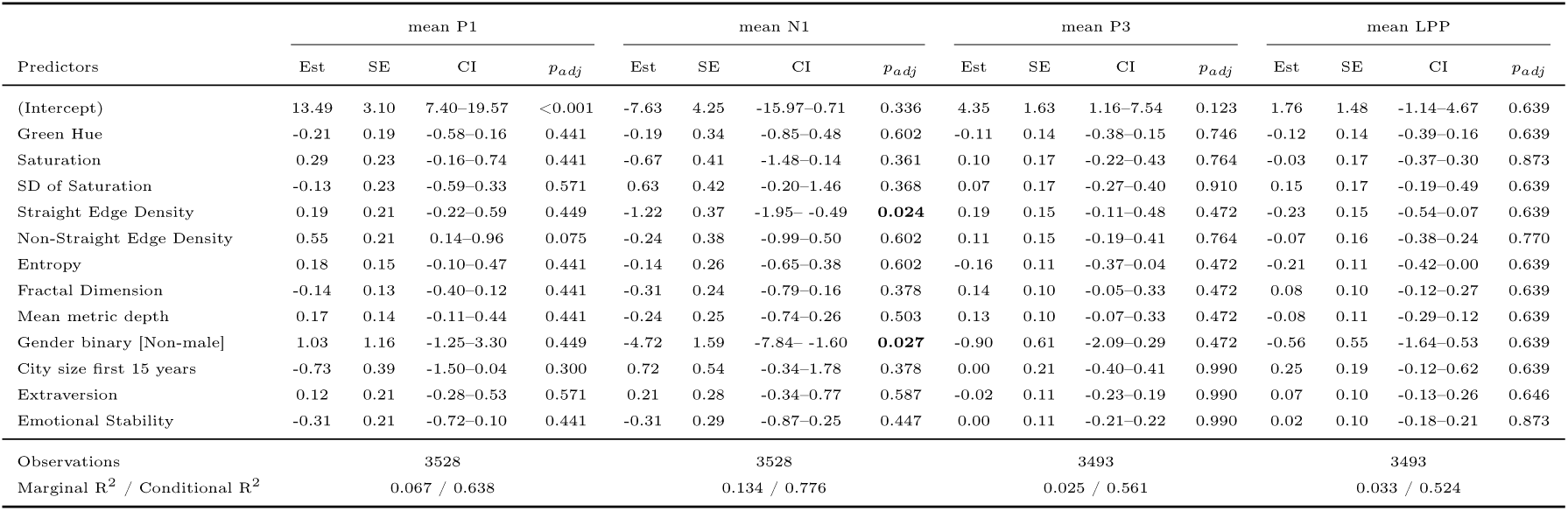
Results of LMMs predicting ERP component amplitudes from LLVFs. *Note: CI are 95% confidence intervals; p-values were adjusted using FDR method; all predictors were z-scaled*.

### 3.3. Associations between ERPs and Ratings

To explore possible connections between ERPs and subjective ratings, indicating an effect of perceptual processing of visual urban stimuli as well as subsequent appraisal processes on subsequent ratings, we calculated nine LMMs with the amplitudes of the four ERP components as predictors and subjective ratings as outcomes. All LMMs contained a random intercept per subject, as well as the same personality characteristics as the previous models as covariates. All p-values were adjusted for multiple hypothesis testing using FDR. For full results, see Tables 7 and 8.

**Table 7:**
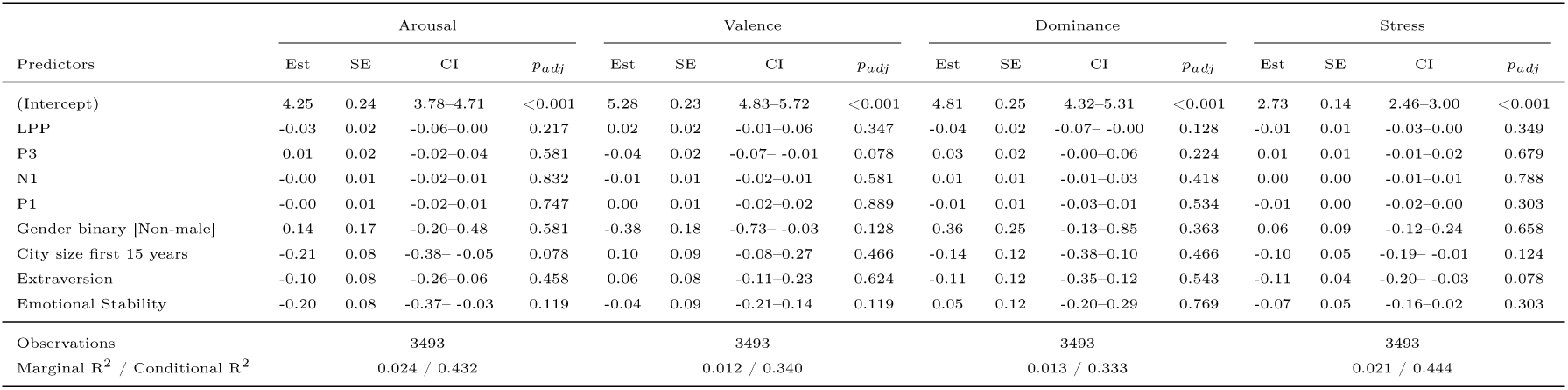
Results of LMMs predicting subjective ratings from ERP component amplitudes (1/2). *Note: CI are 95% confidence intervals; p-values were adjusted using FDR method*.

**Table 8:**
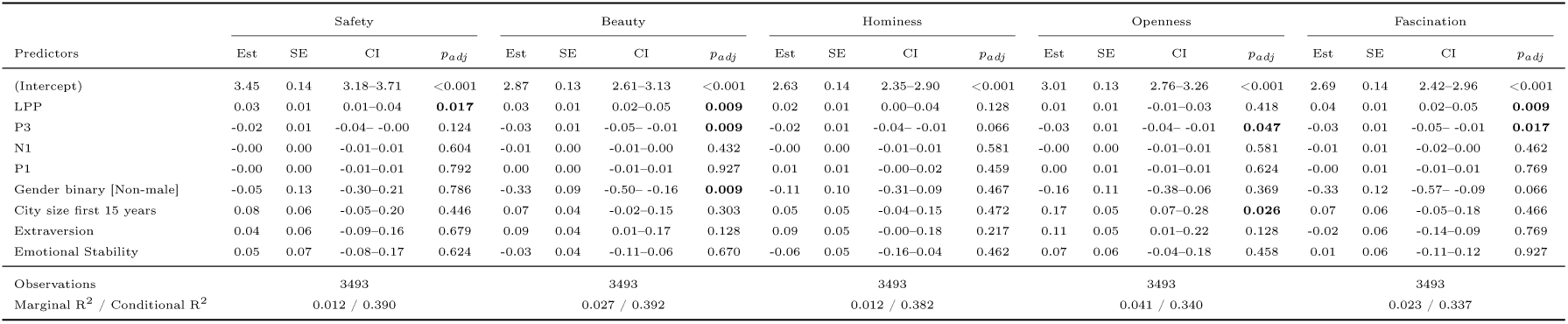
Results of LMMs predicting subjective ratings from ERP component amplitudes (2/2). *Note: CI are 95% confidence intervals; p-values were adjusted using FDR method*.

The ERP components P1 and N1 at occipital electrodes were not significantly associated with subjective ratings (all *p* ≥ 0.303). In contrast, the models revealed a significant effect of the parietal components P3 and LPP on subjective ratings. Increasing P3 amplitudes predicted decreasing beauty ratings (*β* = –0.03, *p* = .009), as well as lower perceived openness (*β* = –0.03, *p* = .047) and fascination (*β* = –0.03, *p* = .017). Conversely, LPP amplitudes were significantly associated with perceived safety (*β* = 0.03, *p* = .017), beauty (*β* = 0.03, *p* = .009), and fascination (*β* = 0.04, *p* = .009), with higher amplitudes preceding higher subjective ratings.

## 4. Discussion

The present study investigated the influence of semantic contents as well as LLVFs of urban images on subjective and neural responses. For a summary of all significant effects reported in this study, see Table 9.

**Table 9:** Summary of significant predictors across all subjective and ERP outcomes. *Note: Arrows indicate significant effects and their direction*

| Predictors | Arousal | Valence | Dominance | Stress | Safety | Beauty | Hominess | Openness | Fascination | P1 | N1 | P3 | LPP |
| --- | --- | --- | --- | --- | --- | --- | --- | --- | --- | --- | --- | --- | --- |
| Greenery/Built Ratio | ↓ | ↑ | ↓ | ↓ |  | ↑ | ↑ | ↑ |  | ↑ |  |  |  |
| Sky | ↑ | ↓ | ↑ | ↑ | ↓ | ↓ | ↓ | ↑ | ↓ |  |  |  |  |
| Car |  |  |  |  |  |  | ↑ |  |  |  |  |  |  |
| Person |  |  |  |  |  |  |  |  |  |  |  |  |  |
| Green Hue |  |  |  |  |  |  |  |  |  |  |  |  |  |
| Saturation |  |  |  |  |  |  |  |  |  |  |  |  |  |
| SD of Saturation |  |  |  |  |  |  |  |  |  |  |  |  |  |
| Straight Edge Density |  |  |  |  |  | ↑ |  |  | ↑ |  | ↑ |  |  |
| Non-Straight Edge Density |  | ↑ | ↓ |  | ↑ | ↑ |  | ↑ | ↑ |  |  |  |  |
| Entropy |  |  |  |  |  |  |  |  |  |  |  |  |  |
| Fractal Dimension |  |  |  |  |  |  |  |  |  |  |  |  |  |
| Mean Metric Depth | ↑ | ↓ | ↑ | ↑ | ↓ | ↓ |  | ↑ | ↑ |  |  |  |  |
| Gender binary [Non-male] |  |  |  |  |  | ↓ |  |  | ↓ |  | ↓ |  |  |
| City size first 15 years | ↓ |  |  | ↓ |  |  |  | ↑ |  |  |  |  |  |
| Extraversion |  |  |  |  |  | ↑ |  |  |  |  |  |  |  |
| Emotional Stability |  |  |  |  |  |  |  |  |  |  |  |  |  |
| Mean P1 |  |  |  |  |  |  |  |  |  |  |  |  |  |
| Mean N1 |  |  |  |  |  |  |  |  |  |  |  |  |  |
| Mean P3 |  |  |  |  |  | ↓ |  | ↓ | ↓ |  |  |  |  |
| Mean LPP |  |  |  |  | ↑ | ↑ |  |  | ↑ |  |  |  |  |

### 4.1. Subjective Ratings of Urban Scenes

Greenery in urban images consistently exhibited beneficial effects across several psychological dimensions. Images with higher greenery/built ratios were associated with lower arousal and stress ratings, more positive valence, stronger feelings of control, and higher perceived beauty and hominess.

In contrast to previous studies reporting a further positive impact of sky area on affective responses to urban scenes (Asgarzadeh et al., 2014; Peng et al., 2025; Rhee et al., 2023), our results predominantly demonstrated a negative impact of increasing sky areas in the displayed scenes. More sky area in an image was associated with higher arousal and stress ratings, lower valence ratings, and reduced feelings of control and safety, as well as lower beauty, hominess, and fascination ratings. This pattern challenges the assumption that openness and visual access to sky are inherently restorative. As would be expected, when considering LLVFs of the images, a strong positive correlation of visible sky area with spatial depth was observed. Like sky area, depth in images showed a similar negative impact on ratings, suggesting that the negative influence of visible sky area in an image is closely related to the spatial openness of an urban scene. This negative impact of sky area on subjective experience might be explainable with the Prospect-Refuge Theory (Appleton, 1975) extended to urban scenes and architecture (Dosen and Ostwald, 2016), as open spaces with extensive sky maximize prospect while minimizing refuge, potentially resulting in feelings of exposure and vulnerability in urban settings (Dosen and Ostwald, 2016).

Perceived openness and sky visibility can evoke feelings of emptiness, being unwelcoming, or unsafe in general (Cui et al., 2023). This is supported by research on spatial enclosure suggesting that excessive enclosure but also excessive openness are detrimental (Fisher-Gewirtzman, 2018). Some stimuli in the present study may have contained disproportionately wide sky views, resulting in depth and exposure exceeding levels of comfort and control. As, contrary to our findings, sky visibility in natural landscapes is often associated with positive responses and aesthetic appreciation (Franěk, 2023), the appraisal of openness appears to be contextdependent. This suggests that the affective influence of openness may depend on surrounding factors. In urban settings, particularly those dominated by built structures and lacking vegetative refuge, the same openness may signal barrenness or insufficient environmental complexity to sustain interest. Additionally, openness in urban environments often coincides with large streets and intersections, which might also play a significant role in their affective and aesthetic appraisal.

### 4.2. Neural Dynamics of Urban Scene Perception

The positive impact of greenery in urban scenes in our study was reflected in a modulation of the visual evoked P1 component approximately 105 ms after scene onset. Higher proportions of natural elements in the presented images elicited significantly increased P1 amplitudes. While in this study, the effect was specific only to the semantic category of greenery but not to related LLVFs, the effect of NSED (which is characteristic of greenery) approached significance. It is thus reasonable to consider that there might also be preferential early visual processing for LLVFs that are characteristic of natural elements, but that the effect did not survive multiple hypothesis correction and a resulting loss of power in the present study design.

These results are theoretically significant, as they replicate previous findings (Grassini et al., 2019) and provide converging evidence for some form of preferential early visual processing of natural environmental features. They may also explain previous behavioral findings which suggest approach biases and faster reaction times in response to natural as compared to urban scenes (Schiebel et al., 2022). This early differentiation may set the stage for subsequent affective processing, though our data revealed no direct connection between early neural responses and subjective appraisal, hinting at a more complex relationship.

The subsequent N1 component, in contrast to the P1, showed selective sensitivity to SED, with higher rectilinearity in an image predicting more negative amplitudes. Straight edges mainly occur in built/man-made structures, suggesting that the observed N1 modulation reflects sensitivity to the geometric properties of scenes. This effect may help explain previous findings (Grassini et al., 2019) which showed that urban scenes elicit stronger N1 responses than natural scenes. This result, contrasting with the earlier P1 component, suggests that different stages of early visual processing might prioritize different aspects of the visually perceived environment. During scene perception, lower-level features that are characteristic of natural elements seem to be preferentially processed before those characteristic of artificial/built elements.

### 4.3. Neural Correlates of Scene Appraisal

Another central finding of our study is the relationship between later ERP components and subjective ratings. While early visual evoked components over the occipital cortex reflected automated processing of natural categories and artificial lower-level features, amplitudes of the P1-N1-complex were not associated with subjective ratings of urban scenes. Conversely, the later P3 and LPP components demonstrated no modulation by environmental features, but a systematic association with subjective ratings.

The observed negative relationship between P3 amplitude and ratings of openness, beauty, and fascination may indicate more effortful attentional processing. Since larger P3 amplitudes are generally thought to reflect greater allocation of attentional resources (Polich, 2012), this pattern could suggest that less pleasant scenes demand increased cognitive effort for processing and evaluation.

A complementary explanation, which might be consistent with this effort-related interpretation, is that positively experienced environments are processed more efficiently, whereas potentially negative or less inviting environments require more extensive accumulation of sensory evidence. In decision-making research, the amplitude of the centroparietal P3 has been shown to increase systematically with the accumulation of sensory evidence (Kelly and O’Connell, 2013; O’Connell et al., 2012). If the P3 amplitude reflects the amount of evidence required to reach a decision threshold, our results may indicate that appraising unpleasant scenes involves more effortful evidence gathering and thus more demanding evaluative processing.

The positive association between LPP and ratings of safety, beauty, and fasci-nation supports interpretations of the component being associated with sustained attention and more elaborate processing (Cuthbert et al., 2000; Schupp et al., 2000). Individuals showing larger LPP amplitudes may have engaged more elaborately in scene evaluation, investing greater attentional resources in aesthetic appraisal.

Together, our results regarding P3 and LPP amplitudes appear to capture complementary but distinct aspects of environmental appraisal. While both components are related to evaluations of aspects of the environment, they likely index different processing stages. The P3 component might reflect earlier, effort-related attentional engagement, while the LPP might reflect later, more elaborative motivational processing. The two components also show partially selective associations with scene attributes, suggesting at least partially independent contributions to the perception of urban environments.

These findings are partially consistent with the evolutionary-constructivist perspective proposed by Koivisto et al. (2022). The modulation of P1 and N1 amplitudes by the proportion of greenery and straight edges suggests that early neural processes are sensitive to the physical properties of the stimuli, which may reflect bottom-up influences. The dissociation between these early sensory components and later evaluative components (P3/LPP) may further indicate that top-down processes contribute to appraisal at a later processing stage. However, the extent to which early and late components are dependent on bottom-up and top-down processes remains unclear from the present data alone. The present results may thus be compatible with a sequential relationship between early stimulus-driven processing, potentially reflecting phylogenetic influences, and later evaluative processes, likely reflecting ontogenetic factors. Future research would be needed to test this interpretation.

In conclusion, we believe the investigated environmental features likely act as building blocks that, together with many other factors, shape experiences of and affective responses to urban environments. While the earlier components P1/N1 directly reflect perceptual responses modulated by these environmental features, later subjective ratings are also influenced by them. There appears to be a clear link between both, although the exact foundational mechanics establishing this link between perception and appraisal remain to be investigated further in future research.

### 4.4. Limited Replication of Effects

A pattern across our findings is the limited number of significant effects for LLVFs and semantic image contents. Among eight extracted LLVFs that have previously been shown to significantly affect our outcome variables (Celikors and Wells, 2022; Ma et al., 2023; Samaan et al., 2024), our results could only replicate the effects of edge density measures and depth, while the effects of green pixels, saturation, brightness, entropy, and fractal dimensions failed to be replicated. Similarly, among semantic categories, only greenery/built ratios and sky area showed robust effects, while the presence of cars and people had no significant influence on any outcome measures. This finding contrasts with our previous study using the same experimental protocol and stimuli (Sander et al., 2024), where these predictors showed small but significant effects. This discrepancy can possibly be explained by originally small effects combined with reduced statistical power in our selected subset of stimuli. Additionally, in the present study, a larger number of models led to stronger multiple hypothesis correction, causing many predictors to fail to achieve statistical significance.

Beyond these considerations, the limited predictor effects likely stem from the visual complexity inherent in ecologically valid images (Sander et al., 2024). In naturalistic urban scenes, environmental features are embedded in multivariate contexts where numerous properties covary, making it difficult to isolate individual feature contributions. Prior research demonstrating the effects of specific features often used controlled and simplified stimuli where the feature of interest was the primary varying dimension. In our ecologically more valid approach, single quantifiable features may be overshadowed by a multitude of other properties. It should also be noted that although several significant effects were detected in this study, the overall variance explained by the fixed effects of environmental features is rather low with marginal *R*^2^ values ranging from 0.064 – 0.221 (*M* = 0.13). This is likely explained in part by the fact that while the stimuli varied in their composition with respect to the environmental features of interest, the images were still generally quite similar. They all represent images of the same city and thus effect sizes are expected to be limited compared to using starkly differing environments (e.g., cities vs. forests).

### 4.5. Limitations and Future Directions

Although our sample size was sufficiently large, it should be noted that we did not assess participants’ color vision capabilities. Therefore, we cannot with certainty say whether all participants perceived color correctly, which might limit the analyses regarding effects of the color composition of the stimulus material. The stimulus material of the present study was selected for ecological validity and captures the actual visual complexity people encounter in urban scenes. This realism, however, also reproduces the inherent relationships and co-occurrences among various environmental features. Such interdependencies make it challenging to isolate the specific contribution of individual factors using simple linear models and global image statistics. Consequently, our analytical approach may have still been overly reductive for the complex stimulus set, and subtle feature effects may have been masked by uncontrolled variability within the scenes. Additionally, the high multicollinearity between semantic predictors and LLVFs further complicates interpretation, reflecting natural co-occurrence patterns that cannot be experimentally disentangled without artificial stimulus manipulation. This limitation constrains causal inference and leaves open whether observed effects reflect truly distinct influences or correlated properties. A related but distinct interpretative constraint applies to the greenery/built ratio itself, as a higher ratio is simultaneously consistent with more vegetation and with fewer built structures, meaning that the reported effects of this predictor cannot be attributed exclusively to the presence of greenery, the absence of built elements, or their combination. Moreover, global image properties such as mean saturation and total sky area may be too coarse to capture what humans actually attend to. Viewers might not experience scenes as uniform averages but selectively attend to salient regions in the stimulus. While semantic segmentation provides an efficient and reproducible method for quantifying the compositional properties of urban scenes, pixellevel classifications of environmental categories (e.g., “greenery”) capture the physical presence of these elements but do not reflect their perceptual salience or affective meaning. Elements assigned to the same semantic category may differ substantially in their restorative or aesthetic impact, and the psychological effects of broadly defined categories are unlikely to be equivalent across subtypes. Beyond these considerations, grouping together environmental categories like buildings, vehicles, plants, etc., disregards nuances that might arise from different details of these objects, e.g., their specific materiality, architectural style, or type. Future work should therefore examine whether and how the precision of segmentation-based approaches can be improved by incorporating sub-categorial distinctions and assess the generalizability of such predictors across different urban contexts. Region-specific analyses might better predict responses (Fernandes and Castelhano, 2021). Future studies jointly analyzing EEG and eye-tracking could provide insights about what aspects of a scene participants focused on and, hence, which image properties were actually processed. Complementary qualitative analyses could help further understand what informs and drives people’s appraisal of their environments. More generally, while the segmentation approach used in this study allowed for a fine-grained quantification of our stimuli, nuanced investigation of the influences of urban features beyond dichotomies, the pretrained model was not fine-tuned on stimulus material comparable to ours. Most categories of interest were detected with satisfactory accuracy, except for the “person” category. Accordingly, the lack of detected effects of people present in the scenes should be interpreted cautiously. Future research should consider more recent state-of-the-art models, potentially fine-tuned on street-level urban imagery comparable to the stimulus material. Furthermore, although specific familiarity with the depicted Berlin environments was not assessed, the statistical models included city size during upbringing as a partial control for general familiarity with urban settings. Specific prior experiences with the shown environments remain a potential uncontrolled source of individual variation, and future studies should examine more directly how subjective experience shapes responses to urban scenes. Moreover, while the experimental protocol would have allowed to investigate a potential impact of semantic pre-stimulus cues on the subjective and physiological responses, the focus of the present study was on the impact of stimulus composition on subjective experiences and the associated event-related responses. Since we averaged 9 responses across all cue types for the analyses presented here, any systematic difference based on specific cues was averaged out and is thus reflected in the overall effects. Future studies will investigate whether a specific cue (e.g., relaxed – stressful) guides attention to specific aspects of the overall stimulus composition and thus might impact subjective and physiological responses.

Lastly, static images might not allow for capturing the dynamic aspects of real-world experiences, and future research should address the issue of complex naturalistic stimuli through complementary approaches. Virtual reality (VR) experiments could create highly realistic but systematically manipulated urban scenes where specific features vary independently, enabling more reliable interpretations while preserving the immersive experience. Alternatively, exploration of fringe-case urban environments and careful selection of pre-quantified scenes might enable the use of naturalistic stimuli by assuring sufficient independence while maintaining variability in predictors. Finally, field studies with mobile EEG could validate laboratory findings in real-world contexts, strongly increasing ecological validity.

### 4.6. Conclusion

The present study investigated subjective and neural responses to urban environments presented as static images. Subjectively, greenery in the scenes consistently promoted positive outcomes while sky and depth showed negative effects, challenging oversimplified assumptions about perceived openness and sky view, and underscoring the importance of context. This study demonstrates that neural responses to urban environments reveal both early categorical differentiation and later evaluative processes. Modulations of the visual evoked P1 by natural greenery suggest preferential early processing of biophilic content before artificial or built aspects of environments, reflected in N1 modulations. Later event-related components likely reflect evaluative properties, with the P3 component potentially reflecting evidence accumulation for the subsequent response, reflecting differences in response thresholds for different aesthetic evaluations, and the LPP indexing sustained engagement with the stimuli. The disconnect between early visual processing and later evaluative processes associated with specific ratings suggests a complex relationship and potential mediating processes between perception and appraisal. The specifics of this relationship remain elusive. To better understand human responses to urban scenes and their specific characteristics, neural, behavioral, and personal data must be integrated to enable fine-grained analysis of perception-related brain activity. This should best be done in the real world to achieve the highest ecological validity possible to serve as the baseline for controlled laboratory replications.

## CRediT authorship contribution statement

**Carolina Zähme:** Writing – original draft, Writing – review & editing, Formal analysis, Data curation, Investigation, Visualization, Software. **Isabelle Sander:** Writing – original draft, Writing – review & editing, Investigation, Formal Analysis, Visualization, Conceptualization, Methodology, Data curation. **Aleksandrs Kose-levs:** Writing – review & editing, Software, Formal Analysis. **Simone Kühn:** Writing – review & editing, Conceptualization. **Klaus Gramann:** Writing – review & editing, Supervision, Methodology, Conceptualization, Validation, Resources, Funding Acquisition.

## Declarations

### Data availability statement

The data analyzed in this study is available in the BIDS format in an OpenNeuro repository (doi:10.18112/openneuro.ds006850.v1.0.0). The full stimulus set used in the experiment, along with the code for experimental implementation, is available at https://github.com/BeMoBIL/urban_appraisal-experiment. All analysis scripts can be found at: https://github.com/BeMoBIL/perception2appraisal_analyses.

### AI declaration statement

Generative AI (ChatGPT-5) has been used only to improve the readability and language of parts of the work.

## Acknowledgments

We thank Timo Berg for his help with implementing the experiment and Kim Aljoscha Bressem for his assistance with data collection.

