## Supplementary materials for "From Perception to Appraisal: Brain Responses to Natural and Built Features in Urban Environments" (from another version of this preprint: version 2, 2026-02-25)

### 1 Subjective Measures & SemSeg Models

model 1: $mean\_rating \sim z\_green\_imperv + z\_sky + z\_car + z\_person + (1|Condition) + (1|Subject)$

model 2: $mean\_rating \sim z\_green\_imperv + z\_sky + z\_car + z\_person + Gender\_binary + City\_15 + extrov + emostab + (1|Condition) + (1|Subject)$

model 3: $mean\_rating \sim z\_green\_imperv + z\_sky + z\_car + z\_person + Gender\_binary + City\_15 + extrov + emostab + (1|Condition) + (1 + z\_green\_imperv|Subject)$

model 4: $mean\_rating \sim z\_green\_imperv + z\_sky + z\_car + z\_person + Gender\_binary + City\_15 + extrov + emostab + (1|Condition) + (1 + z\_green\_imperv + z\_sky|Subject)$

model 5: $mean\_rating \sim z\_green\_imperv + z\_sky + z\_car + z\_person + Gender\_binary + City\_15 + extrov + emostab + (1|Condition) + (1 + z\_green\_imperv + z\_sky + z\_car + z\_person|Subject)$

Table 1: Model comparison for Arousal~SemSeg

| Models | npar | AIC | BIC | logLik | deviance | Chisq | Df | Pr(>Chisq) |
|---|---|---|---|---|---|---|---|---|
| arousal model 1 | 8 | 13633 | 13682 | -6808.5 | 13617 | | | |
| arousal model 2 | 12 | 13626 | 13700 | -6800.8 | 13602 | 15.310 | 4 | 0.00410 |
| **arousal model 3** | 14 | 13572 | 13658 | -6772.0 | 13544 | 57.627 | 2 | 3.066e-13 |
| arousal model 4 | 17 | 13569 | 13674 | -6767.3 | 13535 | 9.452 | 3 | 0.02385 |
| arousal model 5 | 26 | 13566 | 13726 | -6756.9 | 13514 | 20.855 | 9 | 0.01331 |

Table 2: Model fit for Arousal~SemSeg

| Model | R2m | R2c |
|---|---|---|
| arousal model 3 | 0.1284504 | 0.4585143 |

Table 3: Model comparison for Valence~SemSeg

| Models | npar | AIC | BIC | logLik | deviance | Chisq | Df | Pr(>Chisq) |
|---|---|---|---|---|---|---|---|---|
| valence model 1 | 8 | 14018 | 14067 | -7001.0 | 14002 | | | |
| valence model 2 | 12 | 14020 | 14094 | -6998.0 | 13996 | 5.9084 | 4 | 0.2060979 |
| valence model 3 | 14 | 13911 | 13997 | -6941.5 | 13883 | 133.1028 | 2 | < 2.2e-16 |
| **valence model 4** | 17 | 13892 | 13997 | -6928.9 | 13858 | 25.1085 | 3 | 1.465e-05 |
| valence model 5 | 26 | 13878 | 14038 | -6913.0 | 13826 | 31.8051 | 9 | 0.0002152 |

Table 4: Model fit for Valence~SemSeg

| Model | R2m | R2c |
|---|---|---|
| valence model 4 | 0.1036597 | 0.3946762 |

Table 5: Model comparison for Dominance∼SemSeg

| Models | npar | AIC | BIC | logLik | deviance | Chisq | Df | Pr(>Chisq) |
|---|---|---|---|---|---|---|---|---|
| dominance model 1 | 8 | 14300 | 14350 | -7142.2 | 14284 | | | |
| dominance model 2 | 12 | 14304 | 14378 | -7140.2 | 14280 | 3.9539 | 4 | 0.4122833 |
| dominance model 3 | 14 | 14226 | 14312 | -7098.9 | 14198 | 82.5996 | 2 | < 2.2e-16 |
| **dominance model 4** | 17 | 14215 | 14320 | -7090.5 | 14181 | 16.8544 | 3 | 0.0007572 |
| dominance model 5 | 26 | 14200 | 14360 | -7073.8 | 14148 | 33.3180 | 9 | 0.0001175 |

Table 6: Model fit for Dominance∼SemSeg

| Model | R2m | R2c |
|---|---|---|
| dominance model 4 | 0.07192687 | 0.3839105 |

Table 7: Model comparison for Safety∼SemSeg

| Models | npar | AIC | BIC | logLik | deviance | Chisq | Df | Pr(>Chisq) |
|---|---|---|---|---|---|---|---|---|
| safety model 1 | 8 | 9469.8 | 9519.1 | -4726.9 | 9453.8 | | | |
| safety model 2 | 12 | 9474.4 | 9548.4 | -4725.2 | 9450.4 | 3.4001 | 4 | 0.4932 |
| safety model 3 | 14 | 9431.5 | 9517.9 | -4701.8 | 9403.5 | 46.8589 | 2 | 6.679e-11 |
| **safety model 4** | 17 | 9412.6 | 9517.5 | -4689.3 | 9378.6 | 24.9165 | 3 | 1.607e-05 |
| safety model 5 | 26 | 9416.6 | 9577.0 | -4682.3 | 9364.6 | 14.0100 | 9 | 0.1220 |

Table 8: Model fit for Safety∼SemSeg

| Model | R2m | R2c |
|---|---|---|
| safety model 4 | 0.07601526 | 0.430407 |

Table 9: Model comparison for Beauty∼SemSeg

| Models | npar | AIC | BIC | logLik | deviance | Chisq | Df | Pr(>Chisq) |
|---|---|---|---|---|---|---|---|---|
| beauty model 1 | 8 | 10144.3 | 10194 | -5064.2 | 10128.3 | | | |
| beauty model 2 | 12 | 10132.4 | 10206 | -5054.2 | 10108.4 | 19.948 | 4 | 0.0005114 |
| beauty model 3 | 14 | 10016.8 | 10103 | -4994.4 | 9988.8 | 119.555 | 2 | < 2.2e-16 |
| **beauty model 4** | 17 | 9980.0 | 10085 | -4973.0 | 9946.0 | 42.836 | 3 | 2.666e-09 |
| beauty model 5 | 26 | 9948.2 | 10109 | -4948.1 | 9896.2 | 49.746 | 9 | 1.203e-07 |

Table 10: Model fit for Beauty∼SemSeg

| Model | R2m | R2c |
|---|---|---|
| beauty model 4 | 0.1341876 | 0.4496917 |

Table 11: Model comparison for Hominess∼SemSeg

| Models | npar | AIC | BIC | logLik | deviance | Chisq | Df | Pr(>Chisq) |
|---|---|---|---|---|---|---|---|---|
| hominess model 1 | 8 | 10351 | 10400 | -5167.5 | 10335 | | | |
| hominess model 2 | 12 | 10352 | 10426 | -5164.1 | 10328 | 6.6627 | 4 | 0.1548 |
| hominess model 3 | 14 | 10208 | 10294 | -5089.9 | 10180 | 148.3751 | 2 | < 2.2e-16 |
| **hominess model 4** | 17 | 10180 | 10285 | -5072.9 | 10146 | 34.0922 | 3 | 1.894e-07 |
| hominess model 5 | 26 | 10126 | 10286 | -5037.0 | 10074 | 71.7092 | 9 | 7.032e-12 |

Table 12: Model fit for Hominess∼SemSeg

| Model | R2m | R2c |
|---|---|---|
| hominess model 4 | 0.1044532 | 0.4439733 |

Table 13: Model comparison for Openness∼SemSeg

| Models | npar | AIC | BIC | logLik | deviance | Chisq | Df | Pr(>Chisq) |
|---|---|---|---|---|---|---|---|---|
| openness model 1 | 8 | 10009.7 | 10059.1 | -4996.9 | 9993.7 | | | |
| openness model 2 | 12 | 10000.2 | 10074.2 | -4988.1 | 9976.2 | 17.588 | 4 | 0.001485 |
| openness model 3 | 14 | 9944.6 | 10031.0 | -4958.3 | 9916.6 | 59.547 | 2 | 1.174e-13 |
| **openness model 4** | 17 | 9853.9 | 9958.7 | -4909.9 | 9819.9 | 96.757 | 3 | < 2.2e-16 |
| openness model 5 | 26 | 9846.6 | 10006.9 | -4897.3 | 9794.6 | 25.290 | 9 | 0.002666 |

Table 14: Model fit for Openness∼SemSeg

| Model | R2m | R2c |
|---|---|---|
| openness model 4 | 0.1155894 | 0.4029517 |

Table 15: Model comparison for Fascination∼SemSeg

| Models | npar | AIC | BIC | logLik | deviance | Chisq | Df | Pr(>Chisq) |
|---|---|---|---|---|---|---|---|---|
| fascination model 1 | 8 | 10408 | 10457 | -5195.9 | 10392 | | | |
| fascination model 2 | 12 | 10407 | 10481 | -5191.4 | 10383 | 8.9599 | 4 | 0.062110 |
| **fascination model 3** | 14 | 10301 | 10388 | -5136.7 | 10273 | 109.3648 | 2 | < 2.2e-16 |
| fascination model 4 | 17 | 10292 | 10396 | -5128.8 | 10258 | 15.8853 | 3 | 0.001197 |
| fascination model 5 | 26 | 10285 | 10446 | -5116.5 | 10233 | 24.4635 | 9 | 0.003626 |

Table 16: Model fit for Fascination∼SemSeg

| Model | R2m | R2c |
|---|---|---|
| fascination model 3 | 0.06426781 | 0.3806685 |

Table 17: Model comparison for Stress∼SemSeg

| Models | npar | AIC | BIC | logLik | deviance | Chisq | Df | Pr(>Chisq) |
|---|---|---|---|---|---|---|---|---|
| stress model 1 | 8 | 13633 | 13682 | -6808.5 | 13617 | | | |
| stress model 2 | 12 | 13626 | 13700 | -6800.8 | 13602 | 15.310 | 4 | 0.00410 |
| **stress model 3** | 14 | 13572 | 13658 | -6772.0 | 13544 | 57.627 | 2 | 3.066e-13 |
| stress model 4 | 17 | 13569 | 13674 | -6767.3 | 13535 | 9.452 | 3 | 0.02385 |
| stress model 5 | 26 | 13566 | 13726 | -6756.9 | 13514 | 20.855 | 9 | 0.01331 |

Table 18: Model fit for Stress∼SemSeg

| Model | R2m | R2c |
|---|---|---|
| stress model 3 | 0.1284504 | 0.4585143 |

#### 2 ERP measures & SemSeg Models

model 1: $mean\_amplitude \sim z\_green\_imperv + z\_sky + z\_car + z\_person + (1|Condition) + (1|Subject)$

model 2: $mean\_amplitude \sim z\_green\_imperv + z\_sky + z\_car + z\_person + Gender\_binary + City\_15 + extrov + emostab + (1|Condition) + (1|Subject)$

model 3: $mean\_amplitude \sim z\_green\_imperv + z\_sky + z\_car + z\_person + Gender\_binary + City\_15 + extrov + emostab + (1|Condition) + (1 + z\_green\_imperv|Subject)$

model 4: $mean\_amplitude \sim z\_green\_imperv + z\_sky + z\_car + z\_person + Gender\_binary + City\_15 + extrov + emostab + (1|Condition) + (1 + z\_green\_imperv + z\_sky|Subject)$

model 5: $mean\_amplitude \sim z\_green\_imperv + z\_sky + z\_car + z\_person + Gender\_binary + City\_15 + extrov + emostab + (1|Condition) + (1 + z\_green\_imperv + z\_sky + z\_car + z\_person|Subject)$

Table 19: Model comparison for P1~SemSeg

| Models | npar | AIC | BIC | logLik | deviance | Chisq | Df | Pr(>Chisq) |
|---|---|---|---|---|---|---|---|---|
| p1 model 1 | 8 | 19400 | 19449 | -9692.0 | 19384 | | | |
| p1 model 2 | 12 | 19402 | 19476 | -9688.9 | 19378 | 6.1419 | 4 | 0.18880 |
| **p1 model 3** | 14 | 19380 | 19466 | -9675.8 | 19352 | 26.3516 | 2 | 1.896e-06 |
| p1 model 4 | 17 | 19385 | 19490 | -9675.5 | 19351 | 0.5641 | 3 | 0.90460 |
| p1 model 5 | 26 | 19383 | 19544 | -9665.6 | 19331 | 19.7761 | 9 | 0.01935 |

Table 20: Model fit for P1~SemSeg

| Model | R2m | R2c |
|---|---|---|
| p1 model 3 | 0.05335347 | 0.6597734 |

Table 21: Model comparison for N1~SemSeg

| Models | npar | AIC | BIC | logLik | deviance | Chisq | Df | Pr(>Chisq) |
|---|---|---|---|---|---|---|---|---|
| n1 model 1 | 8 19745 19795 -9864.7 | 19729 | | | | | | |
| n1 model 2 | 12 | 19742 | 19816 | -9859.2 | 19718 | 10.9258 | 4 | 0.027411 |
| **n1 model 3** | 14 | 19735 | 19822 | -9853.6 | 19707 | 11.2078 | 2 | 0.003683 |
| n1 model 4 | 17 | 19740 | 19844 | -9852.8 | 19706 | 1.7133 | 3 | 0.633981 |
| n1 model 5 | 26 | 19740 | 19900 | -9843.7 | 19688 | 18.0530 | 9 | 0.034563 |

Table 22: Model fit for N1~SemSeg

| Model | R2m | R2c |
|---|---|---|
| n1 model 3 | 0.1155035 | 0.791748 |

Table 23: Model comparison for P3∼SemSeg

| Models | npar | AIC | BIC | logLik | deviance | Chisq | Df | Pr(>Chisq) |
|---|---|---|---|---|---|---|---|---|
| p3 model 1 | 8 | 15685 | 15735 | -7834.7 | 15669 | | | |
| **p3 model 2** | 12 | 15691 | 15765 | -7833.5 | 15667 | 2.3203 | 4 | 0.6771 |
| p3 model 3 | 14 | 15694 | 15780 | -7832.9 | 15666 | 1.1523 | 2 | 0.5620 |
| p3 model 4 | 17 | 15698 | 15802 | -7831.7 | 15664 | 2.4077 | 3 | 0.4922 |
| p3 model 5 | 26 | 15703 | 15863 | -7825.6 | 15651 | 12.3013 | 9 | 0.1969 |

Table 24: Model fit for P3∼SemSeg

| Model | R2m | R2c |
|---|---|---|
| p3 model 2 | 0.02032965 | 0.5802735 |

Table 25: Model comparison for LPP∼SemSeg

| Models | npar | AIC | BIC | logLik | deviance | Chisq | Df | Pr(>Chisq) |
|---|---|---|---|---|---|---|---|---|
| lpp model 1 | 8 | 15657 | 15707 | -7820.7 | 15641 | | | |
| **lpp model 2** | 12 | 15662 | 15736 | -7819.0 | 15638 | 3.3905 | 4 | 0.4947 |
| lpp model 3 | 14 | 15664 | 15750 | -7817.8 | 15636 | 2.3490 | 2 | 0.3090 |
| lpp model 4 | 17 | 15668 | 15773 | -7817.0 | 15634 | 1.5675 | 3 | 0.6668 |
| lpp model 5 | 26 | 15676 | 15836 | -7812.1 | 15624 | 9.9288 | 9 | 0.3563 |

Table 26: Model fit for LPP∼SemSeg

| Model | R2m | R2c |
|---|---|---|
| lpp model 2 | 0.02593185 | 0.5433793 |

### 3 Subjective Measures & LLVFs

model 1: $mean\_rating \sim z\_green\_mask\_density + z\_Saturation + z\_sdSaturation + z\_StraightEdgeD + z\_NonStraightED + z\_Entropy + z\_FractalD + z_depth\_metric\_mean + (1|Condition) + (1|Subject)$

model 2: $mean\_rating \sim z\_green\_mask\_density + z\_Saturation + z\_sdSaturation + z\_StraightEdgeD + z\_NonStraightED + z\_Entropy + z\_FractalD + z\_depth_metric_mean + Gender\_binary + City\_15 + extrov + emostab + (1|Condition) + (1|Subject)$

model 3: $mean\_rating \sim z\_green\_mask\_density + z\_Saturation + z\_sdSaturation + z\_StraightEdgeD + z\_NonStraightED + z\_Entropy + z\_FractalD + z\_depth\_metric\_mean + Gender\_binary + City\_15 + extrov + emostab + (1|Condition) + (1 + z\_gvHue|Subject)$

model 4: $mean\_rating \sim z\_green\_mask\_density + z\_Saturation + z\_sdSaturation + z\_StraightEdgeD + z\_NonStraightED + z\_Entropy + z\_FractalD + z\_depth\_metric\_mean + Gender\_binary + City\_15 + extrov + emostab + (1|Condition) + (1 + z\_depth\_metric\_mean|Subject)$

model 5: $mean\_rating \sim z\_green\_mask\_density + z\_Saturation + z\_sdSaturation + z\_StraightEdgeD + z\_NonStraightED + z\_Entropy + z\_FractalD + z\_depth\_metric\_mean + Gender\_binary + City\_15 + extrov + emostab + (1|Condition) + (1 + z\_gvHue + z\_depth\_metric\_mean|Subject)$

Table 27: Model comparison for Arousal~LLVFs

| Models | npar | AIC | BIC | logLik | deviance | Chisq | Df | Pr(>Chisq) |
|---|---|---|---|---|---|---|---|---|
| arousal model 1 | 12 | 13618 | 13692 | -6797.3 | 13594 | | | |
| arousal model 2 | 16 | 13611 | 13710 | -6789.6 | 13579 | 15.333 | 4 | 0.004059 |
| arousal model 3 | 18 | 13529 | 13640 | -6746.3 | 13493 | 86.478 | 2 | < 2.2e-16 |
| arousal model 4 | 18 | 13563 | 13674 | -6763.7 | 13527 | 0.000 | 0 | |
| **arousal model 5** | 21 | 13465 | 13595 | -6711.7 | 13423 | 104.046 | 3 | < 2.2e-16 |

Table 28: Model fit for Arousal~LLVFs

| Model | R2m | R2c |
|---|---|---|
| arousal model 5 | 0.1952753 | 0.4871342 |

Table 29: Model comparison for Valence~LLVFs

| Models | npar | AIC | BIC | logLik | deviance | Chisq | Df | Pr(>Chisq) |
|---|---|---|---|---|---|---|---|---|
| valence model 1 | 12 | 14004 | 14078 | -6989.8 | 13980 | | | |
| valence model 2 | 16 | 14006 | 14104 | -6986.8 | 13974 | 5.9185 | 4 | 0.2053 |
| valence model 3 | 18 | 13905 | 14016 | -6934.4 | 13869 | 104.7193 | 2 | <2e-16 |
| valence model 4 | 18 | 13972 | 14084 | -6968.2 | 13936 | 0.0000 | 0 | |
| **valence model 5** | 21 | 13870 | 14000 | -6914.1 | 13828 | 108.3042 | 3 | <2e-16 |

Table 30: Model fit for Valence~LLVFs

| Model | R2m | R2c |
|---|---|---|
| valence model 5 | 0.1483423 | 0.3992565 |

Table 31: Model comparison for Dominance∼LLVFs

| Models | npar | AIC | BIC | logLik | deviance | Chisq | Df | Pr(>Chisq) |
|---|---|---|---|---|---|---|---|---|
| dominance model 1 | 12 | 14296 | 14370 | -7135.8 | 14272 | | | |
| dominance model 2 | 16 | 14300 | 14398 | -7133.8 | 14268 | 3.9574 | 4 | 0.4118 |
| dominance model 3 | 18 | 14224 | 14335 | -7094.1 | 14188 | 79.3556 | 2 | <2e-16 |
| dominance model 4 | 18 | 14246 | 14358 | -7105.2 | 14210 | 0.0000 | 0 | |
| **dominance model 5** | 21 | 14166 | 14296 | -7062.2 | 14124 | 86.1707 | 3 | <2e-16 |

Table 32: Model fit for Dominance∼LLVFs

| Model | R2m | R2c |
|---|---|---|
| dominance model 5 | 0.08737876 | 0.3979389 |

Table 33: Model comparison for Safety∼LLVFs

| Models | npar | AIC | BIC | logLik | deviance | Chisq | Df | Pr(>Chisq) |
|---|---|---|---|---|---|---|---|---|
| safety model 1 | 12 | 9466.4 | 9540.4 | -4721.2 | 9442.4 | | | |
| safety model 2 | 16 | 9471.0 | 9569.7 | -4719.5 | 9439.0 | 3.4031 | 4 | 0.4928 |
| safety model 3 | 18 | 9391.2 | 9502.3 | -4677.6 | 9355.2 | 83.7324 | 2 | <2e-16 |
| safety model 4 | 18 | 9424.6 | 9535.7 | -4694.3 | 9388.6 | 0.0000 | 0 | |
| **safety model 5** | 21 | 9340.7 | 9470.3 | -4649.4 | 9298.7 | 89.9203 | 3 | <2e-16 |

Table 34: Model fit for Safety∼LLVFs

| Model | R2m | R2c |
|---|---|---|
| safety model 5 | 0.09922267 | 0.4516824 |

Table 35: Model comparison for Beauty∼LLVFs

| Models | npar | AIC | BIC | logLik | deviance | Chisq | Df | Pr(>Chisq) |
|---|---|---|---|---|---|---|---|---|
| beauty model 1 | 12 | 10127.6 | 10202 | -5051.8 | 10103.6 | | | |
| beauty model 2 | 16 | 10115.6 | 10214 | -5041.8 | 10083.6 | 19.98 | 4 | 0.0005041 |
| beauty model 3 | 18 | 9955.9 | 10067 | -4959.9 | 9919.9 | 163.72 | 2 | < 2.2e-16 |
| beauty model 4 | 18 | 10072.6 | 10184 | -5018.3 | 10036.6 | 0.00 | 0 | |
| **beauty model 5** | 21 | 9916.1 | 10046 | -4937.1 | 9874.1 | 162.47 | 3 | < 2.2e-16 |

Table 36: Model fit for Beauty∼LLVFs

| Model | R2m | R2c |
|---|---|---|
| beauty model 5 | 0.20238 | 0.4635782 |

Table 37: Model comparison for Hominess∼LLVFs

| Models | npar | AIC | BIC | logLik | deviance | Chisq | Df | Pr(>Chisq) |
|---|---|---|---|---|---|---|---|---|
| hominess model 1 | 12 | 10348 | 10422 | -5161.8 | 10324 | | | |
| hominess model 2 | 16 | 10349 | 10448 | -5158.4 | 10317 | 6.6676 | 4 | 0.1545 |
| hominess model 3 | 18 | 10218 | 10329 | -5091.0 | 10182 | 134.9363 | 2 | <2e-16 |
| hominess model 4 | 18 | 10315 | 10426 | -5139.6 | 10279 | 0.0000 | 0 | |
| **hominess model 5** | 21 | 10182 | 10311 | -5070.0 | 10140 | 139.3384 | 3 | <2e-16 |

Table 38: Model fit for Hominess∼LLVFs

| Model | R2m | R2c |
|---|---|---|
| hominess model 5 | 0.1372554 | 0.4509161 |

Table 39: Model comparison for Openness∼LLVFs

| Models | npar | AIC | BIC | logLik | deviance | Chisq | Df | Pr(>Chisq) |
|---|---|---|---|---|---|---|---|---|
| openness model 1 | 12 | 9972.3 | 10046.4 | -4974.2 | 9948.3 | | | |
| openness model 2 | 16 | 9962.7 | 10061.4 | -4965.3 | 9930.7 | 17.6470 | 4 | 0.001446 |
| openness model 3 | 18 | 9857.2 | 9968.2 | -4910.6 | 9821.2 | 109.5015 | 2 | < 2.2e-16 |
| openness model 4 | 18 | 9852.5 | 9963.6 | -4908.3 | 9816.5 | 4.6416 | 0 | |
| **openness model 5** | 21 | 9748.6 | 9878.1 | -4853.3 | 9706.6 | 109.9365 | 3 | < 2.2e-16 |

Table 40: Model fit for Openness∼LLVFs

| Model | R2m | R2c |
|---|---|---|
| openness model 5 | 0.1665983 | 0.4113985 |

Table 41: Model comparison for Fascination∼LLVFs

| Models | npar | AIC | BIC | logLik | deviance | Chisq | Df | Pr(>Chisq) |
|---|---|---|---|---|---|---|---|---|
| fascination model 1 | 12 | 10393 | 10467 | -5184.6 | 10369 | | | |
| fascination model 2 | 16 | 10392 | 10491 | -5180.1 | 10360 | 8.9756 | 4 | 0.06171 |
| fascination model 3 | 18 | 10276 | 10387 | -5119.8 | 10240 | 120.5565 | 2 | < 2e-16 |
| fascination model 4 | 18 | 10376 | 10487 | -5169.9 | 10340 | 0.0000 | 0 | |
| **fascination model 5** | 21 | 10261 | 10390 | -5109.4 | 10219 | 121.0367 | 3 | < 2e-16 |

Table 42: Model fit for Fascination∼LLVFs

| Model | R2m | R2c |
|---|---|---|
| fascination model 5 | 0.113529 | 0.3861649 |

Table 43: Model comparison for Stress∼LLVFs

| Models | npar | AIC | BIC | logLik | deviance | Chisq | Df | Pr(>Chisq) |
|---|---|---|---|---|---|---|---|---|
| stress model 1 | 12 | 9763.8 | 9837.8 | -4869.9 | 9739.8 | | | |
| stress model 2 | 16 | 9756.6 | 9855.3 | -4862.3 | 9724.6 | 15.1352 | 4 | 0.004429 |
| stress model 3 | 18 | 9726.7 | 9837.8 | -4845.4 | 9690.7 | 33.8728 | 2 | 4.412e-08 |
| stress model 4 | 18 | 9717.3 | 9828.3 | -4840.6 | 9681.3 | 9.4982 | 0 | |
| **stress model 5** | 21 | 9679.0 | 9808.5 | -4818.5 | 9637.0 | 44.2431 | 3 | 1.340e-09 |

Table 44: Model fit for Stress∼LLVFs

| Model | R2m | R2c |
|---|---|---|
| stress model 5 | 0.2182001 | 0.4889557 |

### 4 ERP measures & LLVFs Models

model 1: $mean\_amplitude \sim z\_green\_mask\_density + z\_Saturation + z\_sdSaturation + z\_StraightEdgeD + z\_NonStraightED + z\_Entropy + z\_FractalD + z\_depth\_metric\_mean + (1|Condition) + (1|Subject)$
model 2: $mean\_amplitude \sim z\_green\_mask\_density + z\_Saturation + z\_sdSaturation + z\_StraightEdgeD + z\_NonStraightED + z\_Entropy + z\_FractalD + z\_depth\_metric\_mean + Gender\_binary + City\_15 + extrov + emostab + (1|Condition) + (1|Subject)$
model 3: $mean\_amplitude \sim z\_green\_mask\_density + z\_Saturation + z\_sdSaturation + z\_StraightEdgeD + z\_NonStraightED + z\_Entropy + z\_FractalD + z\_depth\_metric\_mean + Gender\_binary + City\_15 + extrov + emostab + (1|Condition) + (1 + z\_gvHue|Subject)$
model 4: $mean\_amplitude \sim z\_green\_mask\_density + z\_Saturation + z\_sdSaturation + z\_StraightEdgeD + z\_NonStraightED + z\_Entropy + z\_FractalD + z\_depth\_metric\_mean + Gender\_binary + City\_15 + extrov + emostab + (1|Condition) + (1 + z\_depth\_metric\_mean|Subject)$
model 5: $mean\_amplitude \sim z\_green\_mask\_density + z\_Saturation + z\_sdSaturation + z\_StraightEdgeD + z\_NonStraightED + z\_Entropy + z\_FractalD + z\_depth\_metric\_mean + Gender\_binary + City\_15 + extrov + emostab + (1|Condition) + (1 + z\_gvHue + z\_depth\_metric\_mean|Subject)$

Table 45: Model comparison for P1∼LLVFs

| Models | npar | AIC | BIC | logLik | deviance | Chisq | Df | Pr(>Chisq) |
|---|---|---|---|---|---|---|---|---|
| p1 model 1 | 12 | 19405 | 19479 | -9690.6 | 19381 | | | |
| **p1 model 2** | 16 | 19407 | 19506 | -9687.5 | 19375 | 6.1421 | 4 | 0.1888 |
| p1 model 3 | 18 | 19409 | 19520 | -9686.7 | 19373 | 1.5887 | 2 | 0.4519 |
| p1 model 4 | 18 | 19408 | 19519 | -9685.9 | 19372 | 1.5103 | 0 | |
| p1 model 5 | 21 | 19414 | 19543 | -9685.9 | 19372 | 0.0426 | 3 | 0.9977 |

Table 46: Model fit for P1∼LLVFs

| Model | R2m | R2c |
|---|---|---|
| p1 model 2 | 0.06671421 | 0.6377697 |

Table 47: Model comparison for N1∼LLVFs

| Models | npar | AIC | BIC | logLik | deviance | Chisq | Df | Pr(>Chisq) |
|---|---|---|---|---|---|---|---|---|
| n1 model 1 | 12 | 19735 | 19809 | -9855.6 | 19711 | | | |
| **n1 model 2** | 16 | 19732 | 19831 | -9850.1 | 19700 | 10.9297 | 4 | 0.02737 |
| n1 model 3 | 18 | 19730 | 19841 | -9847.1 | 19694 | 6.1461 | 2 | 0.04628 |
| n1 model 4 | 18 | 19728 | 19839 | -9845.9 | 19692 | 2.2966 | 0 | |
| n1 model 5 | 21 | 19724 | 19854 | -9841.3 | 19682 | 9.3075 | 3 | 0.02547 |

Table 48: Model fit for N1∼LLVFs

| Model | R2m | R2c |
|---|---|---|
| n1 model 2 | .1340953 | 0.7763703 |

Table 49: Model comparison for P3∼LLVFs

| Models | npar | AIC | BIC | logLik | deviance | Chisq | Df | Pr(>Chisq) |
|---|---|---|---|---|---|---|---|---|
| p3 model 1 | 12 | 15687 | 15761 | -7831.7 | 15663 | | | |
| **p3 model 2** | 16 | 15693 | 15792 | -7830.6 | 15661 | 2.3201 | 4 | 0.6771 |
| p3 model 3 | 18 | 15695 | 15806 | -7829.6 | 15659 | 1.8678 | 2 | 0.3930 |
| p3 model 4 | 18 | 15694 | 15804 | -7828.8 | 15658 | 1.7388 | 0 | |
| p3 model 5 | 21 | 15696 | 15825 | -7827.0 | 15654 | 3.4667 | 3 | 0.3251 |

Table 50: Model fit for P3∼LLVFs

| Model | R2m | R2c |
|---|---|---|
| p3 model 2 | 0.0252968 | 0.5605299 |

Table 51: Model comparison for LPP∼LLVFs

| Models | npar | AIC | BIC | logLik | deviance | Chisq | Df | Pr(>Chisq) |
|---|---|---|---|---|---|---|---|---|
| lpp model 1 | 12 | 15658 | 15732 | -7817.0 | 15634 | | | |
| **lpp model 2** | 16 | 15663 | 15761 | -7815.4 | 15631 | 3.3900 | 4 | 0.4948 |
| lpp model 3 | 18 | 15666 | 15777 | -7814.9 | 15630 | 0.9632 | 2 | 0.6178 |
| lpp model 4 | 18 | 15664 | 15775 | -7814.1 | 15628 | 1.5599 | 0 | |
| lpp model 5 | 21 | 15669 | 15799 | -7813.6 | 15627 | 0.8832 | 3 | 0.8295 |

Table 52: Model fit for LPP∼LLVFs

| Model | R2m | R2c |
|---|---|---|
| lpp model 2 | 0.03288941 | 0.523568 |
